# Facultative endosymbiosis between cellulolytic protists and methanogenic archaea in the gut of the Formosan termite *Coptotermes formosanus*

**DOI:** 10.1101/2024.05.03.592298

**Authors:** Masayuki Kaneko, Tatsuki Omori, Katsura Igai, Takako Mabuchi, Miho Sakai-Tazawa, Arisa Nishihara, Kumiko Kihara, Tsuyoshi Yoshimura, Moriya Ohkuma, Yuichi Hongoh

**Author notes:** Corresponding author: Yuichi Hongoh, Postal address: 2-12-1 Ookayama, Meguro-ku, Tokyo 152-8550, Japan. Masayuki Kaneko and Tatsuki Omori should be considered joint first author. Passed away in May 2021 (Tsuyoshi Yoshimura).

## Abstract

Anaerobic protists frequently harbour methanogenic archaea, which apparently contribute to the hosts’ fermentative metabolism by consuming excess H_2_. However, the ecological properties of endosymbiotic methanogens remain elusive in many cases. Here we investigated the ecology and genome of the endosymbiotic methanogen of the *Cononympha* protists in the hindgut of the termite *Coptotermes formosanus*. Microscopic and 16S rRNA amplicon sequencing analyses revealed that a single species, designated here ‘*Candidatus* Methanobrevibacter cononymphae’, is associated with both *Cononympha leidyi* and *Cononympha koidzumii* and that its infection rate in *Cononympha* cells varied from 0.0 to 99.8% among termite colonies. Fine-scale network analysis indicated that multiple 16S rRNA sequence variants coexisted within a single host cell and that identical variants were present in both *Cononympha* species and also on the gut wall. Thus, ‘*Ca.* Methanobrevibacter cononymphae’ is a facultative endosymbiont, transmitted vertically with frequent exchanges with the gut environment. Indeed, transmission electron microscopy showed escape or uptake of methanogens from/by a *Cononympha* cell. The genome of ‘*Ca.* Methanobrevibacter cononymphae’ showed features consistent with its facultative lifestyle: i.e., the genome size (2.7 Mbp) comparable to those of free-living relatives; the pseudogenization of the formate dehydrogenase gene *fdhA*, unnecessary within the non-formate-producing host cell; the dependence on abundant acetate in the host cell as an essential carbon source; and the presence of a catalase gene, required for colonization on the microoxic gut wall. Our study revealed a versatile endosymbiosis between the methanogen and protists, which may be a strategy responding to changing conditions in the termite gut.

## INTRODUCTION

Methanogenic archaea, or methanogens, play a key role in the final step of the decomposition of organic matter in various anaerobic environments [1]. In the guts of termites, for example, which are keystone animals in the terrestrial carbon cycle, methanogens accounted for 0–10% of the prokaryotic gut community and possibly produce 1–3% of the global methane [2,3]. Methanogens are present in the guts of diverse termite lineages [4,5] and are considered to contribute to the gut ecosystem as H_2_-sinks by producing methane from H_2_ and CO_2_ [6,7]. Methanogens are found on the hindgut epithelium [4,8,9] or within the cells of certain gut protist species [10]. The protists in the termite gut belong to the phylum *Parabasalia* or the order *Oxymonadida* in the phylum *Preaxostyla* and play major roles in the digestion of lignocellulose [11]. These gut protists generally harbour endo- or ectosymbiotic prokaryotes, which likely contribute to the host nutrition by supplying nitrogenous compounds [11] and/or removing H_2_ [12,13]. In addition to the mutualistic symbionts, commensal or parasitic endosymbiotic bacteria are occasionally present [14].

Endosymbiotic methanogens are observed, for example, in the parabasalid protist *Trichomitopsis termopsidis* in the gut of the termite *Zootermopsis angusticolis* [15,16] and the oxymonad protist *Dinenympha parva* in the gut of the termite *Reticulitermes speratus* [17,18]. It has been hypothesized that endosymbiotic methanogens promote wood decomposition of the protist hosts by consuming excess H_2_ [10], but their detailed physiology and ecology remain unclear. Indeed, although there are numerous examples of endosymbiosis between anaerobic protists and methanogens in various environments, the ecological properties of endosymbiotic methanogens, including whether they are obligate or facultative endosymbionts, are elusive in many cases [19,20].

The Formosan subterranean termite *Coptotermes formosanus* is the most destructive pest of wooden constructions in southern China and Japan and has invaded Hawaii and the southern part of the United States [21]. The termite harbours a relatively simple protistan gut community consisting of five unculturable parabasalid species: *Pseudotrichonympha grassii*, *Holomastigotoides hartmanii, Holomastigotoides minor, Cononympha* [*Spirotrichonympha*] *leidyi*, and *Cononympha koidzumii* [22–24]. Among them, endosymbiotic methanogens are found only in *Cononympha* cells [25,26], which are mainly localized in the posterior part of the hindgut [27,28]. *Cononympha* protists numerically dominate the protistan gut community, occupying 2,200–10,900 out of 3,800–12,900 total protist cells per gut [28]. Transcriptomic analysis of *Cononympha* cells suggested that *Cononympha* hydrolysed cellulose, hemicellulose, pectin, and chitin, contributing to both lignocellulose digestion and nitrogen recycling [24].

In the present study, we examined the ecology and metabolic capacity of the uncultured endosymbiotic methanogen of *Cononympha* species in the gut of *Coptotermes formosanus* using a combination of microscopy, fine-scale phylogenetic analysis, and genome sequence analysis. Here, we show evidence of facultative endosymbiosis by horizontal acquisition of the methanogen and its genome features adapted to the lifestyle. Our study sheds light on another aspect of the multi-layered symbiotic system in the termite gut.

## MATERIALS AND METHODS

### Termite collection and methanogen infection rate

*Coptotermes formosanus* (family Rhinotermitidae) were collected from four prefectures in Japan (Table S1). The termite colonies collected in 2019 or later were kept with their nest logs in plastic containers in a laboratory until use. Colonies collected in 2018 or earlier were kept in laboratories for years fed with red pine chips. The entire guts of three worker termites per colony were removed using sterile forceps, and the gut contents were suspended in sterile solution U [29]. The ratio of methanogen-containing *Cononympha* cells to total *Cononympha* cells in individual guts was examined on the basis of F_420_ autofluorescence under an Olympus BX51 epifluorescence microscope. The procedure of CH_4_ measurement emitted by termites and semi-quantitative PCR amplification of the gene for methyl coenzyme M reductase (McrA) are described in Supplemental Methods.

### Collection of single *Cononympha* cells and sequencing of 18S rRNA genes

Single *Cononympha* cells were collected using a Leica AM6000 micromanipulation system and subjected to whole genome amplification (WGA) using the illustra GenomiPhi V2 kit (GE Healthcare) as described previously [30]. To identify the species of *Cononympha*, the 18S rRNA gene was amplified by PCR using *Cononympha*-specific primers designed in this study (Table S2). The PCR conditions are described in Supplemental Methods. The PCR products were purified, cloned, and sequenced using the Sanger method as described previously [30].

### 16S rRNA amplicon sequencing analysis

The V3–V4 region (ca. 400 bp) of the 16S rRNA gene was amplified by PCR with prokaryote-universal primers, Pro341F and Pro805R (Table S2), using Phusion High-Fidelity DNA Polymerase (New England Biolabs), as described previously [31]. Purification, library preparation, and paired-end sequencing (300 bp × 2) on the MiSeq platform with the MiSeq Reagent Kit v3 (Illumina) were conducted as described previously [31]. The paired-end reads were trimmed, quality filtered, and sorted into amplicon sequence variants (ASVs) using DADA2 v1.6 [32]. The ASVs were classified using SINA v1.2.11 [33] with the SILVA v132 database [34]. ASVs classified as *Eukarya* or undetermined and ASVs of which frequency was < 0.1% of the total reads were discarded from subsequent analyses.

### Phylogenetic analysis of 16S rRNA genes

Near full-length 16S rRNA genes of methanogens were amplified by PCR using Phusion Hi-Fidelity DNA Polymerase. Primers M23F and M1382R broadly targeting methanogens were used for gut wall samples, while primer CfC-M23F adjusted to the sequence of the endosymbiotic methanogen was used as the forward primer for endosymbiotic methanogens (Table S2). The PCR conditions and procedures of sequencing and alignment are described in Supplemental Methods. A maximum-likelihood tree was constructed using IQ-TREE v1.6.12 [35] with the TVM+F+I+G4 nucleotide substitution model selected by ModelFinder implemented in IQ-TREE. Network construction of sequence variants (SVs) was conducted using the PopART program depending on Qt v4.8.5 with TCS algorithms [36].

### Fluorescence in situ hybridization (FISH) and transmission electron microscopy (TEM)

To discriminate between *Cononympha leidyi* and *C. koidzumii* cells, oligonucleotide probes specific to each 18S rRNA sequence were designed using ARB [37] (Table S2). FISH was performed as described previously [38] with hybridization at 55°C for 2 h. Observations were conducted under the Olympus BX51 epifluorescence microscope. TEM was performed as described previously [39] using an H-7500 transmission electron microscope (Hitachi).

### Genome sequencing

Single *Cononympha* cells were collected as described above, washed several times in droplets of solution U, and transferred to solution U containing 0.1% Tween 20 (Nacalai Tesque). Prokaryotic cells that leaked out from the ruptured host cell were collected with a glass capillary attached to the Leica AM6000 micromanipulation system and subjected to WGA as described above. Nine WGA samples of single *Cononympha* cells were prepared. These were subjected to paired-end sequencing on the MiSeq platform. After assembling and binning as below, one sample was selected for deeper paired-end sequencing and mate-pair sequencing on MiSeq, and long-read sequencing on the MinION platform (Oxford Nanopore Technologies), based on the estimated genome completeness of the target methanogen. The procedures for library preparation and sequencing are described in Supplemental Methods.

### Genome assembly

The MiSeq paired-end reads were trimmed, quality filtered, and assembled into contigs using SPAdes v3.9 [40]. The contigs were binned using a combination of MyCC [41] and Contig Annotation Tool (CAT) [42]. The results were evaluated for the selection of the best sample. The MinION reads, after being quality trimmed, were assembled together with the quality-trimmed MiSeq paired-end and mate-pair reads into contigs using SPAdes v3.15.2 with the ‘hybrid-assembly’ and ‘meta’ mode [40]. The assembled contigs ≥500 bp were binned using a combination of MyCC, CAT, RAST (http://rast.nmpdr.org/rast.cgi), and BLASTn searches. The bin affiliated with *Methanobrevibacter* was further subjected to scaffolding using LINKS v2.0.1 [43] and gap-closing using TGS-GapCloser [44]. The quality-trimmed MiSeq and MinION reads were mapped onto the gap-closed bin using BBMap v38.96 and Minimap2 v2.24-r1122 [45]. The reads mapped onto the bin were then reassembled using SPAdes v3.15.2 with the ‘hybrid-assembly’ and ‘sc’ mode. Reassembled contigs ≥1,000 bp were binned as described above. Details of quality filtering are described in Supplemental Methods.

### Gene annotation

Finding and functional annotation of genes were performed using a combination of DFAST [46] and BLASTp searches of the NCBI non-redundant (nr) protein database. CRISPR-Cas systems were identified using CRISPRCasFinder [47]. Assignment of COG (clusters of orthologous genes) categories [48] was conducted by RPS-BLAST v2.6.0+ searches of the NCBI Conserved Domain Database v3.16, and the results were manually curated. Pseudogenes were manually identified as described previously [49]. Metabolic pathways were inferred using the KEGG automatic annotation server (KAAS) [50] and KEGG Mapper [51].

### Phylogenomics

To construct a phylogenomic tree, we retrieved genome sequences of *Methanobacteriaceae* as references from GTDB r202 [52] and metagenome-assembled genomes (MAGs) assigned to *Methanobacteriaceae* recently obtained from termite guts [5,53]. Gene prediction, extraction, and alignment are described in Supplemental Methods. A maximum-likelihood tree was constructed using IQ-TREE v1.6.12 with the LG+F+R5 amino acid substitution model selected by ModelFinder. The robustness of the tree topology was evaluated using 1,000 ultrafast bootstrap resamplings. Average nucleotide identity (ANI) and average amino acid identity (AAI) were calculated using ANI/AAI-Matrix [54].

### Comparative genome analysis

The relative abundance of genes assigned to COG functional categories was calculated for genome sequences with >80% estimated completeness (Table S3). Principal component analysis was performed after a central log-ratio transformation using a script (https://github.com/dkato2021/COGplot.git). To compare genome contents with identical criterion, the identification of intact genes or pseudogenes was automatically performed using Pseudofinder v1.1.0 [55] and the NCBI Prokaryotic Genome Annotation Pipeline (PGAP) [48].

## RESULTS

### Discovery of *Cononympha* cells lacking endosymbiotic methanogens

During our preliminary observations of protists in the guts of *Coptotermes formosanus* by epifluorescence microscopy, we unexpectedly found that, in certain *Coptotermes formosanus* colonies, no *Cononympha* cells emitted F_420_ autofluorescence as a characteristic signal of the presence of endosymbiotic methanogens, unlike in other *Coptotermes formosanus* colonies (Fig. 1A–D) or previous reports [25,26,28]. To confirm the absence of methanogens in *Cononympha* cells in such termite colonies, we performed amplicon sequencing analysis of the 16S rRNA V3–V4 region for prokaryotic microbiota associated with individual *Cononympha* cells. A total of 30 single *Cononympha* cells collected from two *Coptotermes formosanus* colonies (O2020a and O2020b in Table S1), in which almost all *Cononympha* cells were without F_420_ autofluorescence, were examined. For comparison, we also conducted the same experiment using 28 single *Cononympha* cells from two colonies (K2019b and K2019c in Table S1), in which almost all *Cononympha* cells exhibited F_420_ autofluorescence. No 16S rRNA sequences of methanogens were obtained from the former 30 *Cononympha* cells, whereas sequences of *Methanobrevibacter* were recovered from all of the latter 28 *Cononympha* cells (Fig. S1).

**Fig. 1.**
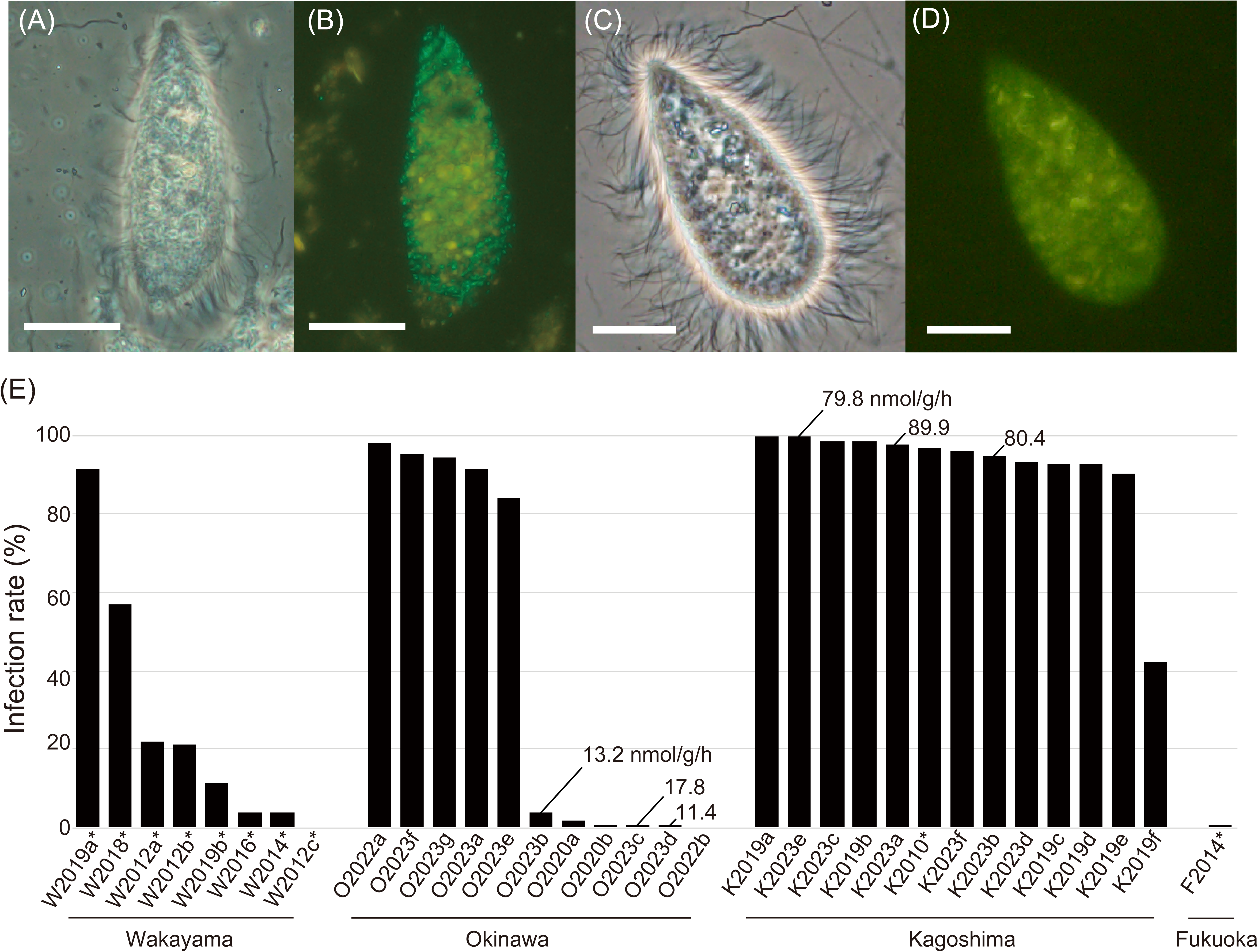
Detection of endosymbiotic methanogens of *Cononympha* protists in the hindgut of *Coptotermes formosanus*. (A, C) Phase-contrast images of *Cononympha* cells. (B, D) Epifluorescent images. Endosymbiotic methanogens are visible with their greenish F_420_ autofluorescence in panel B, whereas no signal was detected in panel D. Amorphous yellow is the autofluorescence of wood particles. Bars indicate 10 µm. (E) Ratio of methanogen-containing *Cononympha* cells to total *Cononympha* cells in individual termite guts across 33 termite colonies. Values indicate methane emission rates (nmol^-1^ g termite^-1^ h^-1^) averaged between biological replicates. Asterisks attached to termite colony IDs indicate laboratory-reared termite colonies.

### Infection rate of methanogens in *Cononympha* cells and methane emission rate

We examined the infection rate of methanogens in *Cononympha* cells for 33 colonies of *Coptotermes formosanus* collected from four prefectures in Japan (Table S1), based on the detection of F_420_ autofluorescence. The infection rate greatly varied from 0 to 99.8% depending on the termite colonies (Fig. 1E; Tables S1 and S4). This difference was irrespective of natural or laboratory-reared termite colonies and prefectures of the sampling sites (Fig. 1E; Tables S1 and S4).

To test whether the presence or absence of the endosymbiotic methanogens of *Cononympha* is linked to the methane emission rate from the termite hosts, we chose three *Coptotermes formosanus* colonies of the high and low endosymbiotic methanogen-infection rate types, respectively (Fig. 1E; Table S1; Supplementary Methods). The results showed that termites from the colonies of the ‘high infection rate’ type emitted 4.5–7.8 times more methane than those from the ‘low infection rate’ type colonies (Fig. 1E). Semi-quantitative PCR targeting the *mcrA* gene in the entire gut microbiota generated results consistent with the methane emission rates (Fig. S2; Supplementary Methods). No *mcrA* amplification was detected in any of the four tested *Coptotermes formosanus* colonies of the ‘low infection rate’ type under the PCR condition, where *mcrA* amplicons were detected in all six tested colonies of the ‘high infection rate’ type (Fig. S2; Table S1).

Because the hindgut wall of *Coptotermes formosanus* can be another habitat of methanogens [56], we examined F_420_ autofluorescence signals on gut wall fragments from termite colonies of both ‘high infection rate’ and ‘low infection rate’ types. We observed dense colonization of rod-shaped methanogens on fragments of the hindgut wall from two out of seven colonies of the ‘high infection rate’ type, whereas almost no signals were detected in termites from four colonies of the ‘low infection rate’ type (Fig. S3; Table S1).

### Fine-scale phylogenetic composition and host specificity of endosymbiotic methanogens

In the above 16S rRNA amplicon sequencing analysis (Fig. S1), multiple ASVs belonging to *Methanobrevibacter* were obtained from each *Cononympha* cell (Fig. S4). Nine closely related ASVs were obtained in total, showing only 1–4 mismatches out of 386 bp. Among them, the ASV-3 sequence was identical to the corresponding region of clone SlMeN10 (AB360373), which was derived from *Cononympha* cells in a *Coptotermes formosanus* gut [26]. Since the genome of the endosymbiotic methanogen contains only a single rRNA operon as described below, these multiple ASVs were not derived from intragenomic variation. These results indicated that multiple strains of a single *Methanobrevibacter* species frequently coexisted within a single *Cononympha* cell.

Next, we examined whether both *Cononympha* species, i.e., *C. leidyi* and *C. koidzumii*, can harbour methanogens. We obtained the 18S rRNA gene sequences of single *Cononympha* cells used in the above 16S rRNA amplicon analysis (Fig. S1) and found that both *Cononympha* species housed methanogens in colonies of the ‘high infection rate’ type (Fig. S5; Table S5). Conversely, in colonies of the ‘low infection rate’ type, both *Cononympha* species lacked methanogens (Fig. S5; Table S5). We observed that both *Cononympha* species, visually discriminated by FISH specifically targeting 18S rRNA of each species, harboured *Methanobrevibacter*-like rods (Fig. S6).

We further investigated whether the lineages of the endosymbiotic methanogens are distinct, even though closely related, between the two *Cononympha* species or not. In addition, we examined the phylogenetic relationship between the endosymbionts and the methanogens on the gut wall. To examine these with higher resolution, we obtained near full-length 16S rRNA gene sequences of methanogens by PCR amplification from 14 single *Cononympha leidyi* cells, three single *C. koidzumii* cells, and seven fragments of the hindgut wall (Fig. 2; Tables S1 and S5). Three out of four 16S rRNA gene SVs of methanogens associated with *C. koidzumii* cells were also obtained from *C. leidyi* cells. In addition, we obtained 16S rRNA gene amplicons of methanogens from three of the seven gut wall fragments and sequenced 24 clones in total, all of which showed 99– 100% sequence similarities to SVs of the endosymbionts (Fig. 2). These SVs from the gut wall or *Cononympha* cells formed clusters that were delineated mainly by the host *Coptotermes formosanus* colonies (Fig. 2); thus, the sequence variations of these methanogens were strongly related to the host termite colony and not to the *Cononympha* species.

**Fig. 2.**
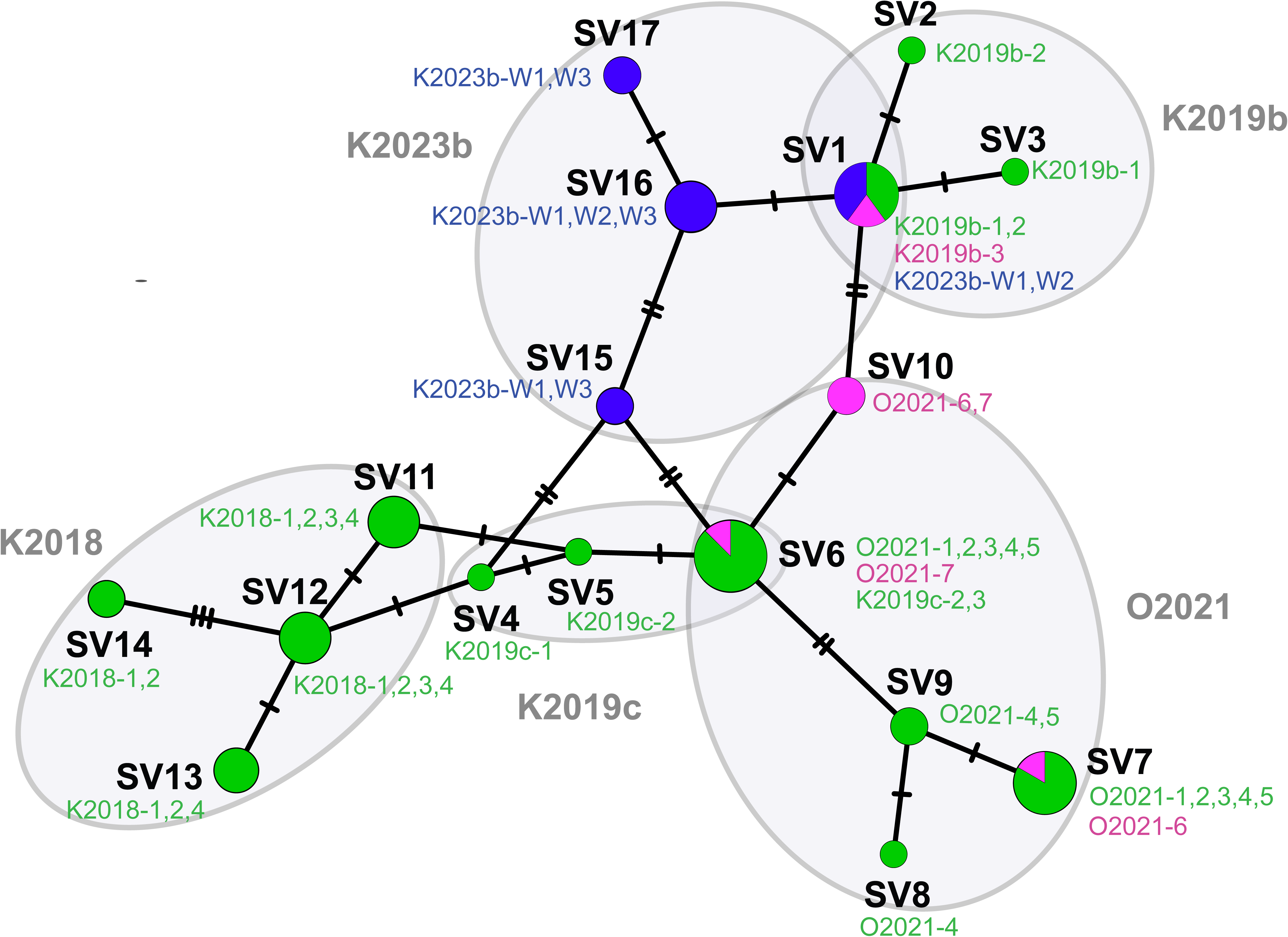
Network analysis of 16S rRNA gene sequence variants (SVs) of methanogens obtained from *Cononympha leidyi* (green), *Cononympha koidzumii* (magenta), and gut wall (blue). Unambiguously aligned 1,312 nucleotide sites were used. The circle size is equivalent to the number of samples of single *Cononympha* cells or gut wall fragments, from which the corresponding SVs were detected. Slashes on branches indicate the number of nucleotide substitutions. The SVs were clustered by termite colonies, and their colony IDs and *Cononympha* cell IDs are shown (see Tables S1 and S5).

These data together indicated that the endosymbiosis between the *Cononympha* protists and the methanogens is facultative and that the two *Cononympha* species can horizontally acquire the same *Methanobrevibacter* species. Furthermore, the spatial distribution of the methanogen most likely extends to the gut wall of the termite host. The shape and size of the methanogens on the gut wall (Fig. S3A) and within *Cononympha* cells were similar. Both methanogens were straight rods, and the size was 1.1–1.8 µm by 0.4–0.5 µm (n = 50) in the former while it was 1.3–1.8 µm by 0.4–0.5 µm (n = 50) in the latter.

### TEM of endosymbiotic methanogens

The facultative nature of this endosymbiosis was further corroborated by TEM. An electron micrograph showed that electron-dense, rod-shaped prokaryotic cells, morphologically indistinguishable, were present not only within a *Cononympha* cell but also on its surface (Fig. 3 and S7A–D). They appeared to be in the process of phago- or exocytosis by the *Cononympha* cell (Fig. 3 and S7A). These prokaryotic cells morphologically resembled cells of cultured *Methanobrevibacter* species [8,9,57]. The intracellular ones were surrounded by a host membrane (Fig. 3 and S7A–D), and some were localized proximal to putative hydrogenosomes (Fig. 3B and S7C). Intracellular *Methanobrevibacter*-like cells apparently during fission were observed (Fig. S7D); they likely proliferate within the host *Cononympha* cell. We hereafter designated this endosymbiotic methanogen as ‘*Candidatus* Methanobrevibacter cononymphae’ (abbreviated as *Mbv. cononymphae*), and the species description is presented below.

**Fig. 3.**
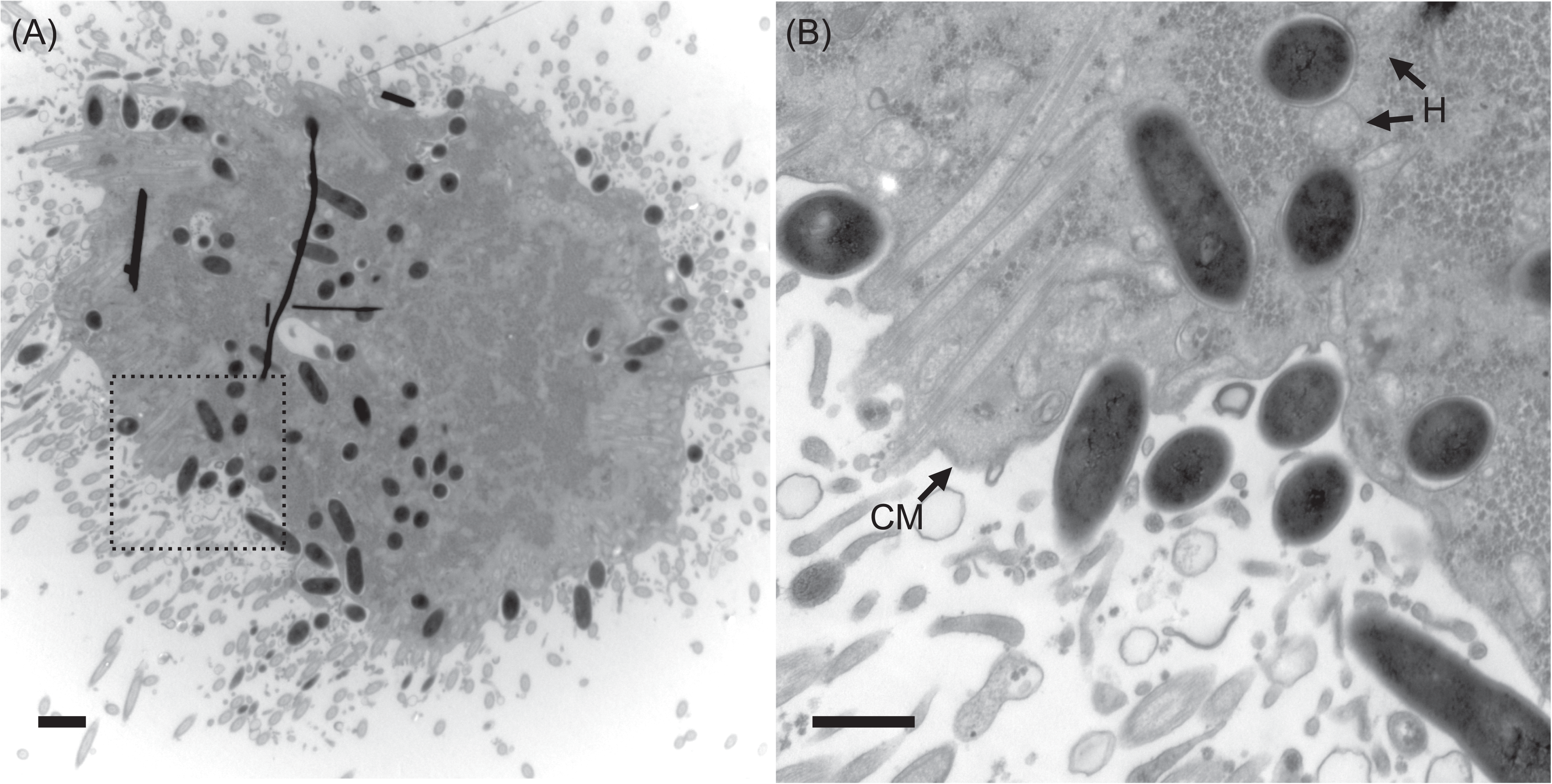
Transmission electron microscopy of a *Cononympha* cell. (A) Section of a *Cononympha* cell. Several black lines are artefacts. (B) Magnified view of the boxed region in panel (A). Electron-dense *Methanobrevibacter*-like cells were observed within the host cytoplasm and also on the host cell surface. H: examples of putative hydrogenosomes; CM: cytoplasmic membrane of *Cononympha*. Bars = 1 µm (A); 500 nm (B).

### Phylogenetic position of *Mbv. cononymphae* based on 16S rRNA gene

Based on the near full-length 16S rRNA gene, *Mbv. cononymphae* was placed within a clade exclusively comprising uncultured clones derived from termite guts (Fig. S8). Within this clade, *Mbv. cononymphae* was closely related to clones obtained from the protists *Microjoenia* sp. and *D. parva* in the gut of *R. speratus* [17]. This clade further formed a monophyletic cluster with termite-gut-derived sequences, including cultured species inhabiting the gut epithelium of the protist-dependent termite *Reticulitermes flavipes*, i.e., *Methanobrevibacter curvatus* [8] and *Methanobrevibacter filiformis* [9]. Endosymbiotic *Methanobrevibacter* housed by an anaerobic ciliate, *Nyctotherus ovalis*, in the hindgut of the cockroach *Blaptica dubia* [58] and that housed by an aquatic anaerobic ciliate, *Trimyema compressum* [20], were more distantly related (Fig. S8).

### General genome features of *Mbv. cononymphae*

We obtained a draft genome sequence of *Mbv. cononymphae* from a single *Cononympha* cell. The genome consisted of 245 contigs with 100% and 98.6% completeness estimated using CheckM [59] and CheckM2 [60], respectively (Table 1). Although only a single rRNA operon was identified in this genome as in many other *Methanobrevibacter* species (Table 1), we recovered four near full-length 16S rRNA gene SVs showing 3–5 base mismatches by PCR amplification from the same DNA sample. Thus, this genome sequence was derived from multiple, at least four, genomovars and here designated as the composite genome ‘CfCl-M3’.

**Table 1.** General genome features of ‘*Ca.* Methanobrevibacter cononymphae’ and.

| species | '*Ca. Mbv. cononymphae*'<br>CfCl-M3 | *Mbv. curvatus*<br>(GCA_0016392<br>95) | *Mbv. filiformis*<br>(GCA_0016392<br>65) | *Mbv. cuticularis*<br>(GCA_0016392<br>85) | NOE<br>(GCA_0033156<br>55) |
| --- | --- | --- | --- | --- | --- |
| Habitat | Intracellular/<br>gut wall | gut wall | gut wall | gut wall | intracellular |
| Total length (Mb) | 2.71 (2.67<sup>d</sup>) | 2.41 | 2.61 | 2.61 | 1.91 |
| Contig | 244 | 232 | 295 | 169 | 70 |
| Completeness<sup>a</sup><br>(%) | 98.6 | 98.2 | 97.1 | 99.7 | 99.3 |
| G+C (%) | 27.2 | 25.7 | 27.0 | 26.8 | 25.3 |
| CDS<sup>b</sup> | 2,319 (1,834) | 1,883 | 1,985 | 1,869 | 2,175 |
| Coding density<br>(%)<sup>b</sup> | 65.8 (59.7) | 68.5 | 68.5 | 66.2 | 68.3 |
| rRNA | 3 | 3 | 3 | 3 | 3 |
| tRNA | 28 | 31 | 30 | 32 | 33 |
| Pseudogene<sup>c</sup> | 29/174 (148) | 11/86 | 9/83 | 15/117 | 95/209 (187) |

<sup>a</sup> Estimated using CheckM2 v.1.0.1.
<sup>b</sup> Identified using Prodigal v2.6.3. Manually curated values are shown in parentheses.
<sup>c</sup> Identified using Pgap/Pseudofinder with those within 1,000 bp region from both contig ends being excluded. The number was manually curated for CfCl-M3 in this study and previously for NOE [57] and shown in parenthesis.
<sup>d</sup> Total length with duplicated CDSs being excluded.

The total contig length was 2,709,899 bp, and 1,834 protein-coding sequences (CDSs) were predicted, excluding 90 CDSs truncated by contig ends and 203 CDSs <100 amino acid sequences that were automatically predicted but with no significant hits in BLASTp searches of the NCBI nr protein database. At least, two CRISPR-Cas systems were identified with a total of 282 spacer sequences (Tables S6–8). Most spacers showed no sequence similarity to those in CRISPRCasdb [61]. The number of pseudogenes was 148 after manual inspection (Table 1), and 14 were assigned to category [V] (defence mechanism) of COGs, including DNA restriction-modification systems and *cas* genes (Fig. S9; Table S6). Finding of duplicated regions and the frequency of single nucleotide polymorphisms in the CfCl-M3 composite genome are described in Supplemental Methods, Supplemental Results, and Table S9.

### Phylogenomics and comparative genome analysis

A maximum-likelihood tree based on concatenated amino acid sequences of conserved single-copy genes (Fig. 4; Table S3) was basically congruent with the 16S rRNA gene tree (Fig. S8). *Mbv. cononymphae* belonged to a monophyletic cluster consisting exclusively of MAGs and two *Methanobrevibacter* isolates obtained from the guts of protist-dependent termites. These genomes shared at least 75% ANI and 60% AAI (Fig. S10). This clade was sister to a clade comprising genome sequences from protist-dependent/independent termite guts or other environments (Fig. 4).

**Fig. 4.**
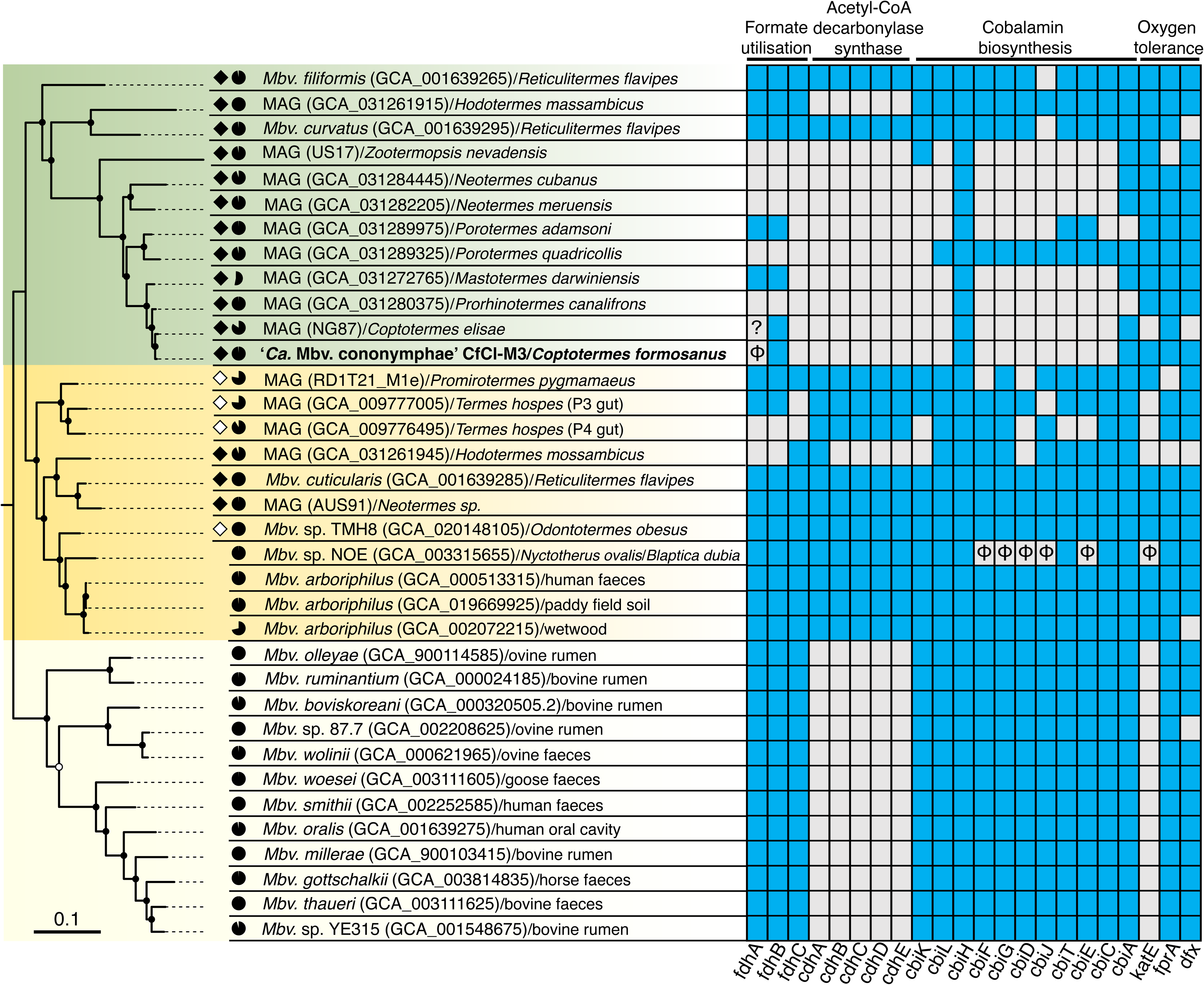
Phylogenetic position and characteristic genetic features of ‘*Ca.* Methanobrevibacter cononymphae’. Maximum likelihood tree was constructed based on 42 concatenated single-copy genes using the LG+F+R5 amino acid substitution model. *Methanobacterium veterum* MK4 and *Methanobacterium bryantii* M.o.H. were used as outgroups (Table S3). Ultrafast bootstrap values of 85–95% and ≥95% are indicated with open and closed circles, respectively. Genomes derived from protist-dependent and protist-independent termite species are highlighted in filled and unfilled diamonds, respectively. The genome completeness estimated using CheckM is shown by pie charts. The presence or absence of genes for formate dehydrogenase FdhAB, formate transporter FdhC, acetyl-CoA decarbonylase/synthase Cdh, cobalamin biosynthesis Cbi, and oxygen tolerance (catalase KatE, F_420_H_2_ oxidase FprA, and desulfoferredoxin Dfx) are linked to the phylogenetic tree. Existing and absent genes are indicated in blue and grey, respectively. Pseudogenes are indicated with ‘ϕ’. The *fdhA* gene of MAG_NG87 is at a contig end and truncated; therefore, it is impossible to specify whether it is intact or pseudogenized.

Genome sequences derived from termite guts in these two clades (Fig. 4) tended to possess more genes assigned to the COG categories [L] (replication, recombination and repair) and [V] (defence mechanisms) (Fig. S11). Among the genes, those related to DNA methylation and drug resistance were especially abundant in termite-gut derived genomes (Fig. S12). No clear differences in the number and ratio of genes assigned to respective COG categories were observed among the five *Methanobrevibacter* species in Table 1, except for a much smaller number and ratio of genes assigned to [V] (defence mechanism) in *Methanobrevibacter* sp. NOE (Fig. S13), which is likely an obligate endosymbiont [57].

The number of pseudogenes of *Mbv. cononymphae* was approximately 1.5–2.1 times larger than those of the gut-wall dwelling *Mbv. curvatus*, *Mbv. filiformis*, and *Methanobrevibacter cuticularis*, but smaller (ca. 80%) than that of *Methanobrevibacter* sp. NOE (Table 1). The distribution pattern of pseudogenes among COG categories indicated that pseudogenes were characteristically accumulated in *Mbv. cononymphae* in categories [L] (replication, recombination, and repair), [V] (defence mechanism), and [X] (mobilome; prophages, transposons) (Fig. S14). No unique genes were identified in *Mbv. cononymphae* among COG-assigned 911 CDSs when compared to other known *Methanobrevibacter* species (Fig. S15; Table S10). Details are described in Supplemental Results.

### Predicted metabolism of *Mbv. cononymphae*

All genes for enzymes required for energy conservation through methanogenesis were present in *Mbv. cononymphae* (Fig. 5). Similar to other *Methanobrevibacter* species, it likely produces methane from H_2_ and CO_2_. On the other hand, the formate dehydrogenase subunit alpha gene *fdhA* is pseudogenized, and the gene for formate transporter (*fdhC*) was missing (Fig. S16). Intact *fdhA* and *fdhC* genes were not found in sequence reads outside the assembled genome or in the other eight samples that were not selected for deeper sequencing analysis. In addition, genes for the acetyl-CoA decarbonylase/synthase (ACDS) multienzyme complex were absent; thus, *Mbv. cononymphae* apparently requires acetate to produce acetyl-CoA. The cobalamin biosynthetic pathway was absent, and instead, genes for the cobalamin/siderophore transporter FepBCD were identified (Table S6). These characteristics, i.e., loss or absence of genes for Fdh, ACDS, and cobalamin biosynthesis, were shared by most MAGs derived from the guts of protist-dependent termite species, but not shared by the gut wall-dwelling species *Mbv. curvatus*, *Mbv. filiformis*, and *Mbv. cuticularis* [8,9] (Fig. 4).

**Fig. 5.**
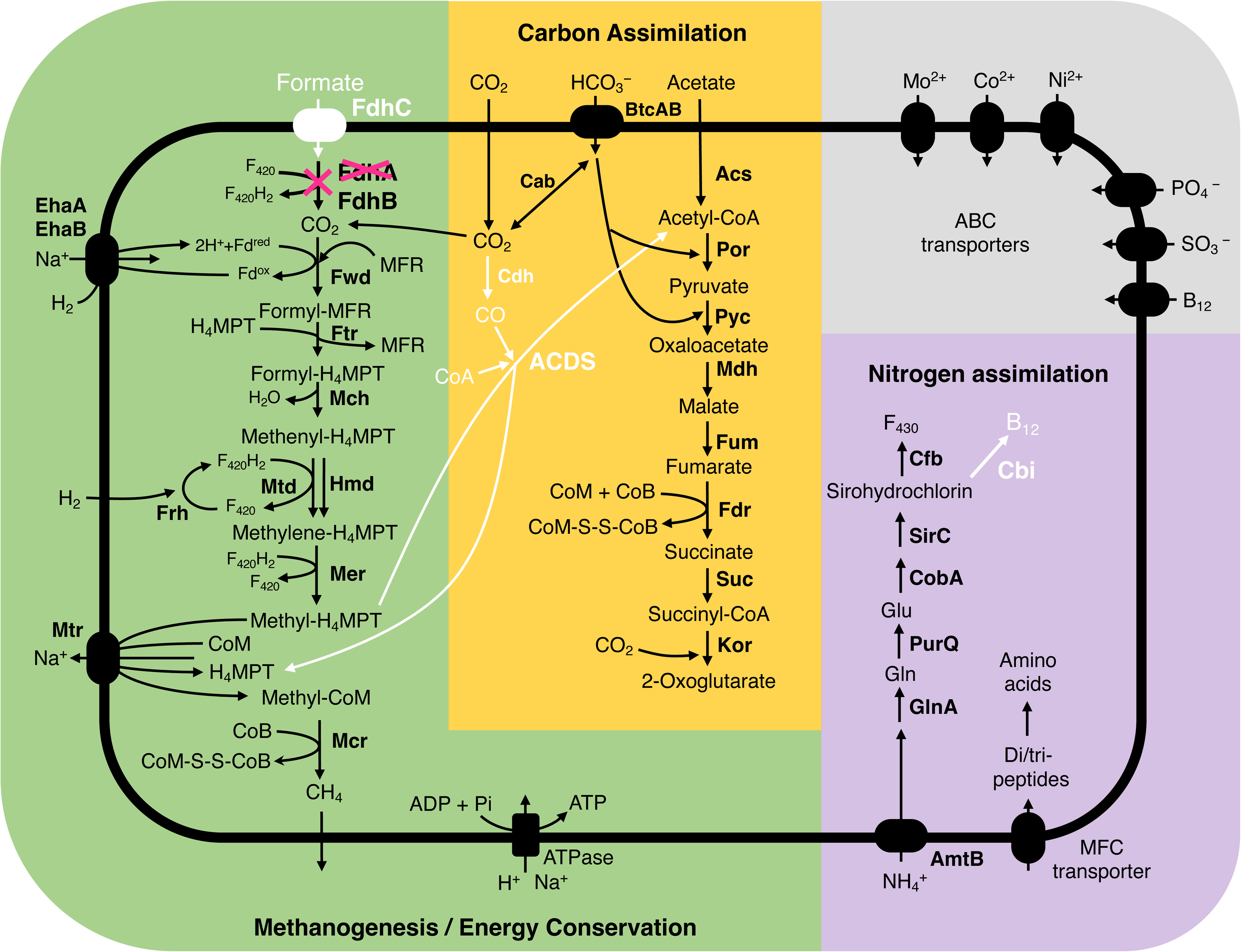
The predicted metabolic pathway of ‘*Ca.* Methanobrevibacter cononymphae’. Genes or pathways characteristically absent in *Mbv. cononymphae* are shown in white. The gene for formate dehydrogenase subunit alpha (*fdhA*) is pseudogenized.

*Mbv. cononymphae* possesses genes for ammonium transporter AmtB and glutamine synthetase GlnA. The genome also encoded a major facilitator superfamily transporter for di- and tripeptides, which may be other nitrogen sources (Fig. 5). The amino acid and cofactor synthesis capabilities were similar to those of other *Methanobrevibacter* species (Table S11). Genes for proteins involved in oxygen tolerance, including catalase KatE, were identified (Fig. 4), and details are described in Supplemental Results. No genes related to motility were found.

## DISCUSSION

This study unveiled the ecology and genomic features of the endosymbiotic methanogen *Mbv. cononymphae* harboured by two *Cononympha* species in the gut of *Coptotermes formosanus*. A series of evidence obtained by a combination of F_420_ autofluorescence detection, small subunit (SSU) rRNA gene sequence analysis, and TEM, clearly indicated that their association is facultative and that the methanogens are most likely transmitted in ‘mixed mode’ [62], i.e., both vertically and horizontally among *Cononympha* cells across the two *Cononympha* species. In addition, our data suggest that *Mbv. cononymphae* colonizes the hindgut wall. This facultative lifestyle is consistent with its genomic features such as the genome size comparable to those of free-living relatives, the number of pseudogenes intermediate between free-living relatives and the putatively obligate endosymbiont NOE of the anaerobic ciliate *N. ovalis* [58] (Table 1), and the presence of a catalase gene (CfCl-M3_0439, Table S6).

Catalase activity was previously observed in *Mbv. filiformis*, *Mbv. curvatus*, and *Mbv. cuticularis*, all of which attach to the microoxic hindgut wall of *R. flavipes* [8,9]. The catalase gene *katE* was commonly found in the clade containing *Mbv. cononymphae* and its sister clade; however, it is pseudogenized in *Methanobrevibacter* sp. NOE (Fig. 4). This is possibly because NOE, unlike *Mbv. cononymphae*, has become an obligate endosymbiont that rarely encounters a high concentration of oxygen.

Assuming that *Cononympha* cells produce H_2_, CO_2_, and acetate as the main fermentation products but do not produce formate, as observed in other cellulolytic parabasalids [63,64] and predicted based on the transcriptome of *Cononympha* [24] (see Supplemental Methods), the loss of *fdhC* and pseudogenization of *fdhA* (Fig. S16) are likely attributable to endosymbiosis. In the hindgut of termites, the amount of formate is generally limited, but still appreciable amounts were detected [7,65]. Indeed, the *fdh* operon is apparently intact in the genomes of the gut wall-dwelling species *Mbv. filiformis*, *Mbv. curvatus*, and *Mbv. cuticularis* (Fig. S16), even though these isolates exhibited only weak or no growth on formate [8,9]. It is conceivable that the reduced opportunity and/or requirement to use formate has allowed the loss of functional *fdh* genes in *Mbv. cononymphae*. The lack of ACDS in *Mbv. cononymphae* may also be related to endosymbiosis. The expectedly high concentration of acetate within the host protist cell may have allowed the endosymbiont to depend solely on acetate to produce acetyl-CoA (Fig. 5). However, as the concentration of acetate is also high in the termite hindgut [7, 65], other factors may be involved in the absence of ACDS.

The absence of the functional biosynthetic pathway of cobalamin, which is essential in methanogenesis [66] (Fig. 5), was previously reported also in the endosymbiotic methanogens *Methanobrevibacter* sp. NOE and *Methanocorpusculum* sp. MCE [58]. Thus, its absence and dependence on a transporter may be a common trait in endosymbiotic methanogens. These characteristics, i.e., the absence of functional *fdh*, ACDS genes, cobalamin biosynthetic pathway, and the presence of the catalase gene were common in most MAGs from the guts of protist-dependent termites (Fig. 4). This implies that these methanogens possibly have a lifestyle similar to that of *Mbv. cononymphae*, i.e., facultative endosymbiosis, although their localizations are unidentified. Among them, MAG_NG87 from *Coptotermes elisae* [53] and MAG_GCA031280375 from *Prorhinotermes canalifrons* [5] showed high ANIs, 98.8% and 96.2%, respectively, with *Mbv. cononymphae*, and both termite genera harbour *Cononympha* species [67]; these MAGs are most likely of endosymbionts of *Cononympha* with similar lifestyles.

It is widely believed that hydrogenotrophic methanogens critically contribute to the fermentative metabolism of H_2_-producing anaerobic protist hosts as syntrophic mutualists [19]. Indeed, retarded growth of anaerobic ciliates that had lost their endosymbiotic methanogens was previously observed [68,69]. In particular, removal of ‘*Candidatus* Methanoregula pelomyxae’, an endosymbiont of *Pelomyxa schiedti*, led to the death of the anaerobic amoeba host [70]. In the case of termite-gut protists, Messer and Lee (1989) [16] reported that *Trichomitopsis termopsidis* treated with 2-bromoethanesulfonate (BES) inhibiting methanogenesis was viable in the gut of *Z. angusticollis*. However, in an experiment using cultured *Trichomitopsis termopsidis,* the growth of the protist was severely retarded when methanogenesis was suppressed by BES treatment [64]. Its growth was recovered when fed with a more favourable nutrient source in addition to cellulose, i.e., autoclaved cells of a specific bacterial strain [64]. This reminded other reports that anaerobic ciliates during successive cultures under optimal conditions tended to lose their endosymbiotic methanogens, whereas ciliates under food-limiting conditions or at low temperatures kept housing methanogens [71,72]. These previous results imply that the loss of *Mbv. cononymphae* may be related to the nutritional condition of the *Cononympha* host. It is unclear whether *Mbv. cononymphae* contributes to the nutrition of the *Cononympha* host, not only by consuming H_2_, but also by supplementing nitrogenous compounds as suggested in several cases of obligate endosymbioses between termite-gut protists and bacteria [12,49,73–75].

Although *Mbv. cononymphae* can also likely colonize the gut wall, the methane emission rate shown in Fig. 1E indicated that the absence of the endosymbiont was not compensated by methanogens outside *Cononympha* cells. Large differences in the methane emission rate among colonies were previously observed in other termite species, i.e., *R. speratus* [2]*, N. sugioi* [*koshunensis*] [56], and *Cryptotermes secundus* [2,7]. Thus, the facultative association of methanogens not only with gut protists but also with termite hosts may be not rare phenomena. Although what factors determine the infection rate of methanogens in gut protists and their total abundance in the termite gut remain unclear, methanogens might play a role in adjusting the fermentation process of the protist hosts and the whole termite gut ecosystem in response to changing nutritional conditions with their versatile localization and abundance.

### Description of ‘*Candidatus* Methanobrevibacter cononymphae’ sp. nov

*Methanobrevibacter cononymphae* (co.no.nym’phae. N.L. fem. n. *Cononymphae*, of *Cononympha,* a genus of flagellated protist hosts). The archaea are straight rods with dimensions of 1.3–1.8 µm (mean ± SD, 1.58 ± 0.1; *n* = 50) by 0.4–0.5 µm (0.4 ± 0.0). The archaea are non-motile and facultatively colonize the cytoplasm of *Cononympha leidyi* and *Cononympha koidzumii* in the gut of *Coptotermes formosanus*. The archaea occasionally colonize the hindgut wall of *Coptotermes formosanus*. The assignment is based on the 16S rRNA gene (LC802674). The draft genome sequence CfCl-M3 showed 96.2% ANI to *Methanovirga procula,* which has been named under SeqCode on the basis of a MAG (GCA_031280375) obtained from the gut of *Prorhinotermes canalifrons* [5]. Since the genus *Methanobrevibacter* is highly divergent, Protasov et al (2023) have proposed division of *Methanobrevibacter* into at least nine genera, including *Methanovirga*, based on the genomic phylogeny and distance under SeqCode [5, 76].

## Supporting information

Supplemental Tables_S1-S11_rev

## DATA AVAILABILITY

The 16S rRNA amplicon and metagenome sequence data generated in this study have been deposited in DDBJ under the accession numbers DRR534530–534599 (BioProject: PRJDB17265). Accession numbers for the CfCl-M3 genome are BAABUN010000001–BAABUN010000245. Representative near full-length SSU rRNA gene sequences will appear under the accession numbers LC802720–61 (18S rRNA) and LC802666–82 (16S rRNA).

## AUTHOR CONTRIBUTIONS

Y.H. and M.O. conceived the study and provided equipment and reagents; Y.H., K.K., and T.Y. designed the study; Y.H. K.K., T.Y., M.K, T.O, and M.S.-T. collected termites; T.O., M.K., M.S.-T., and K.I. performed experiments, M.K., T.O., and Y.H. analysed and interpreted the data; T.M. performed TEM; A.N., M.K, and K.I. measured methane; M.K., K.I., and Y.H. wrote the paper with input from the other authors.

## ACKNOWLEDGEMENTS

We are grateful to Hirokazu Kuwahara for supporting the collection of protist cells and genome sequencing, to Takumi Murakami for handling sequence data, and to Katsuhiro Aminaka for helping with micromanipulation. We thank Yukihiro Kinjo, Osamu Kitade, Shuji Itakura, Satoko Noda, Wakako Omura, Masao Oya, Kazuki Takahashi, Gaku Tokuda, and Akinori Yamada for their cooperation in termite collection. We also thank Gillian Gile for her helpful advice on the morphological discrimination of *Cononympha* species, Satoshi Hattori for his advice on the metabolism of *Methanobrevibacter*, and Naoya Shinzato for unpublished information on endosymbiotic methanogens of *Trimyema compressum*. We thank Michiru Shimizu, Jun-ichi Inoue, and Masahiro Yuki for assisting with Sanger sequencing, which was performed in the RIKEN BSI. Sanger sequencing was also performed in the Biomaterial Analysis Center in the Tokyo Institute of Technology. TEM was assisted by Keiko Ikeda in the latter facility.

## COMPETING INTERSTS

The authors declare no competing interests.

