## Supplementary material for "Facultative endosymbiosis between cellulolytic protists and methanogenic archaea in the gut of the Formosan termite *Coptotermes formosanus*": Supplemenal Text

**SUPPLEMENTAL TEXT**

**SUPPLEMENTAL METHODS**

**Measurement of CH_4_**

Worker termites of *Coptotermes formosanus* from colonies within one week after collection were used (Table S1). Four days before the measurements, termites were transferred from their nest wood to a Petri dish containing cellulose powder (Nacalai Tesque) kneaded with double-distilled water at a ratio of 4:7. The termites were then transferred to a Petri dish with moistened filter paper a day before the measurements. These procedures were required to minimize the effect of food among termite colonies and to easily handle the termites for subsequent gas chromatography. From 30 to 50 termites were placed into 5-mL amber glass vials (net volume 10.2 mL) containing new moistened filter paper. The vials were then sealed with butyl rubber stoppers. The CH_4_ concentration in the vials was measured at 2, 4, and 5 h using a gas chromatograph with a GC-2014 flame ionization detector (Shimadzu) and a Porapak N 80/100 column (GL Sciences) at 100°C. The carrier was nitrogen gas flowing at 25 mL min^-1^. To estimate any methane background generated from non-biological production, a negative control without termites was prepared.

**Semi-quantitative PCR and sequencing of *mcrA***

From 10 to 15 worker termites per colony were used for DNA extraction from the entire gut as described previously [1]. DNA concentration was measured using the Qubit dsDNA HS Assay Kit (Invitrogen) and adjusted to an equal amount among samples. PCR targeting the *mcrA* gene, a marker gene for methanogens, was performed using specific primers (Table S2) and Takara Ex Premier DNA Polymerase (Takara Bio) as follows: 1 min initial denaturation at 94°C, 25 cycles of denaturation (10 s at 98°C), annealing (15 s at 55°C), extension (30 s at 68°C) and a final 4 min extension at 68°C. The PCR products separated by agarose gel electrophoresis were stained with ethidium bromide, and the fluorescence intensities were quantified using Image Lab 5.0 software (Bio-Rad).

To identify the origin of *mcrA*, we increased the PCR cycles to 30 and purified the products using the MonoFas DNA Purification Kit (ANIMOS). After cloning the products using the TOPO TA Cloning Kit for Sequencing (Invitrogen), Sanger sequencing was performed as described previously [2].

**Sequencing of 18S rRNA genes**

The 18S rRNA gene was amplified by PCR using *Cononympha*-specific primers, Con-F and Con-R, designed in this study (Table S2). PCR was performed using Takara *Ex Taq* DNA Polymerase (Takara Bio) as follows: 1 min initial denaturation at 94°C, 30 cycles of denaturation (20 s at 94°C), annealing (30 s at 71.3°C), extension (90 s at 72°C) and a final 5 min extension at 72°C.

**Phylogenetic analysis of 16S rRNA genes**

Near full-length 16S rRNA genes of methanogens in *Cononympha* cells and those attaching to the gut wall were amplified by PCR using Phusion Hi-Fidelity DNA Polymerase (New England Biolabs). The PCR conditions for endosymbiotic methanogens were as follows: 30 s initial denaturation at 98°C, 30 cycles of denaturation (10 s at 98°C), annealing (30 s at 60°C), extension (90 s at 72°C) and a final extension for 4 min at 72°C. For gut-wall methanogens, the annealing temperature was 50°C. The PCR products were purified, cloned, and sequenced using the Sanger method as described by Igai et al. (2022) [2]. The sequences obtained were aligned with reference sequences retrieved from the SILVA v132 and NCBI non-redundant (nr) nucleotide databases using MAFFT v7.490 [3]. The alignment was trimmed using trimAl v1.2 rev59 [4].

**Genome sequencing**

Sequencing libraries were prepared using the QIAseq FX DNA Library kit (Qiagen), and paired-end sequencing (300 bp × 2) was conducted on the MiSeq platform. Among the nine samples, one was selected on the basis of the estimated genome completeness of the target methanogen after assembling and binning, and the sample was subjected to the second-round WGA using the GenomiPhi HY DNA Amplification Kit (GE Healthcare) as described previously [5]. Sequencing libraries for the sample were prepared using the TruSeq DNA PCR-Free Library Prep Kit and the Nextera Mate Pair Library Preparation Kit (Illumina). Paired-end and mate-pair sequencing analyses were conducted on the MiSeq platform as described above.

In addition, a long-read sequencing library was prepared from the same WGA sample. The sample was subjected to debranching and single-strand DNA digestion using EquiPhi29 DNA Polymerase (ThermoFisher) and S1 nuclease (Takara Bio), respectively. The resulting DNA was fragmented using Covaris g-TUBE and separated by agarose gel electrophoresis. Fragments of 3–10 kb were excised and purified using the Zymoclean Large Fragment DNA Recovery Kit (Zymo Research). A library was prepared using the SQK-LSK109 Ligation Sequencing Kit (Oxford Nanopore Technologies), and sequencing was performed using an R10.3 Spot-On Flow cell (FLO-MIN109) on the MinION platform.

**Genome assembly**

Trimming and quality filtering of MiSeq reads were conducted using Cutadapt [6], NxTrim [7], and PRINSEQ [8]. For the MinION reads, base calling was performed using guppy v3.2.10 (Oxford Nanopore Technologies), and adapter removal, quality trimming, and chimeric read removal were performed using Porechop v0.2.4 [9], Nanofilt v2.7.1 [10], and yacrd v0.5.1 [11], respectively.

**Single nucleotide polymorphism (SNP) analysis**

SNPs and indels were calculated using HaplotypeCaller implemented in GATK 4.2.0.0 [12]. Only MiSeq reads with mapping quality ≥30 were used as described previously [13].

**Phylogenomics**

Genes were predicted using Prodigal v2.6.3 [14], and 43 single-copy gene markers used by Parks et al. [15] were extracted using the IdentifyHMM program (https://github.com/edgraham/PhylogenomicsWorkflow). Only genome sequences with at least 22 of the 43 marker genes were used for the phylogenomic analysis (Table S3). Amino acid sequences were aligned using MAFFT v7.470, concatenated, and trimmed using trimAl v1.4 rev22.

**Prediction of *Cononympha* metabolism**

To predict the metabolism of the protist host, a previous transcriptome dataset of *Cononympha* species [16] was retrieved and analysed as follows. First, transcripts less than 1 TPM (transcripts per million) and those derived from prokaryotes, predicted using CAT, were excluded. Then, transcripts mapped onto the target genome sequence using Minimap2 v2.24-r1122 [17] were excluded. Finally, metabolic pathways were inferred using KAAS [18] and KEGG Mapper [19].

**SUPPLEMENTAL RESULTS**

**Origin of *mcrA* from *Coptotermes formosanus* guts**

The *mcrA* sequences were obtained from one whole-gut sample derived from the O2022a colony (‘high infection rate type’) and also one from the O2022b colony (‘low infection-rate type’). The closest sequences were identified using BLASTx searches of the NCBI nr database.

Of 19 sequences obtained from the ‘high infection rate type’ colony, 14 shared 100% amino acid sequence identity with the *mcrA* gene (CfCl-M3_0356) of ‘*Ca*. Methanobrevibacter cononymphae’. The remaining five showed 98.6% amino acid sequence similarity to an uncultured *Methanobrevibacter* clone derived from a biogas reactor (AFL70135). In the ‘low infection rate type’ sample, three of six successfully obtained *mcrA* sequences showed 97.8% amino acid sequence similarity to an uncultured *Methanobrevibacter* clone derived from a biogas reactor (AFL70134). The remaining three showed the highest amino acid sequence similarity (90.7%) to an uncultured *Methanomassiliicoccales* clone from the gut of the termite *Macrotermes michaelseni* (AFU08097). Because *Methanomassiliicoccales* do not emit F_420_ autofluorescence [20], it is possible that *Methanomassiliicoccales* may inhabit the gut of *Coptotermes formosanus* without being detected by epifluorescence microscopy and accounted for the appreciable amounts (11.4–17.8 nmol/g/h) of methane emitted from the termites from the ‘low infection rate type’ colonies (Fig. 1E).

**Duplicated regions and SNPs**

Of 2,319 CDSs automatically identified using Prodigal v2.6.3, 119 CDSs were clustered into 42 groups with ≥99% sequence identity (Table S9). These CDS groups may reflect the presence of redundant regions that were derived from multiple, coexisting genomovars and were not assembled together possibly because of genome rearrangement. Indeed, the value 42 outnumbered those of isolated *Methanobrevibacter* strains or *Methanobrevibacter* sp. NOE (Table S9). When these redundant regions (~2% of the draft genome sequence) were excluded, the total contig length was ca*.* 2.67 Mb. The frequency of single nucleotide polymorphisms and indels within the population in the single host cell was 3.13 per kilobase pair in total, which was higher than 0.77 and 0.80 in the obligate endosymbionts ‘*Ca*. Adiutrix trichonymphae’ Adiu2019 and ‘*Ca*. Endomicrobium trichonymphae’ Rs-D17, respectively [13].

**Shared and unique genes in *Mbv. cononymphae* among *Methanobrevibacter***

*Mbv. cononymphae* shared 856 CDSs out of COG-assigned 911 CDSs with either *Mbv. curvatus*, *Mbv. filiformis*, or *Mbv. cuticularis*, and 55 were unique (Fig. S15). Of these 55 CDSs, ten were not found in *Methanobrevibacter* genomes obtained from environments other than termite guts, and only one, a pseudogene of AAA family ATPase, was not found in any *Methanobrevibacter* genome (Table S10).

**Genes involved in oxygen tolerance**

In addition to the catalase gene *katE*, the *Mbv. cononymphae* genome encoded genes for F_420_H_2_ oxidase FprA (CfCl-M3_1464), rubrerythrin FprB (CfCl-M3_1465), desulfoferredoxin (superoxide reductase-like protein; CfCl-M3_1820), rubredoxin domain-containing rubrerythrin (CfCl-M3_1821), and rubredoxin (CfCl-M3_623, 624, 1734) (Table S5). Combinations of these proteins can detoxify O_2_, H_2_O_2_, or O_2_^-^ to some extent [21,22], and unlike *katE*, these genes were identified in the genomes of most *Methanobrevibacter* species, including NOE (Fig. 4).

**REFERENCES TO SUPPLEMENTAL TEXT**
