## Supplementary Figures_S1-S16 for "Facultative endosymbiosis between cellulolytic protists and methanogenic archaea in the gut of the Formosan termite *Coptotermes formosanus*"

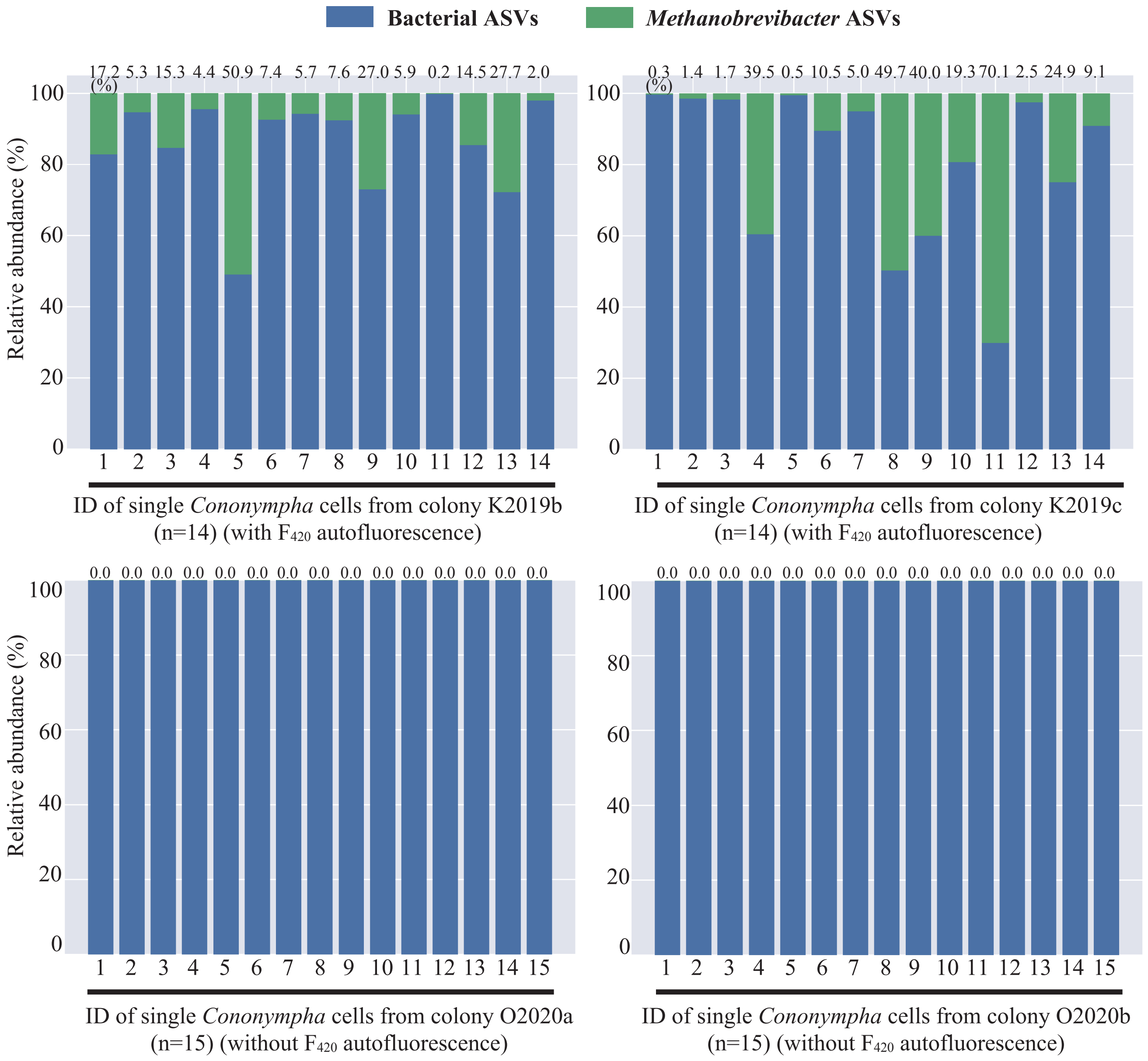


**Fig. S1.** Taxonomic composition of prokaryotes associated with single *Cononympha* cells based on 16S rRNA V3–V4 amplicon sequences. Four or five single *Cononympha* cells per worker individual were subjected to whole genome amplification and then to PCR amplification and sequencing on the MiSeq platform (12,497–73,225 read pairs per sample after being quality trimmed). The numerical values on the bar graph indicate the ratio of amplicon sequence variants assigned to *Methanobrevibacter* among all prokaryotic sequence reads. See Table S1 for information on the four termite colonies used here.

(A)

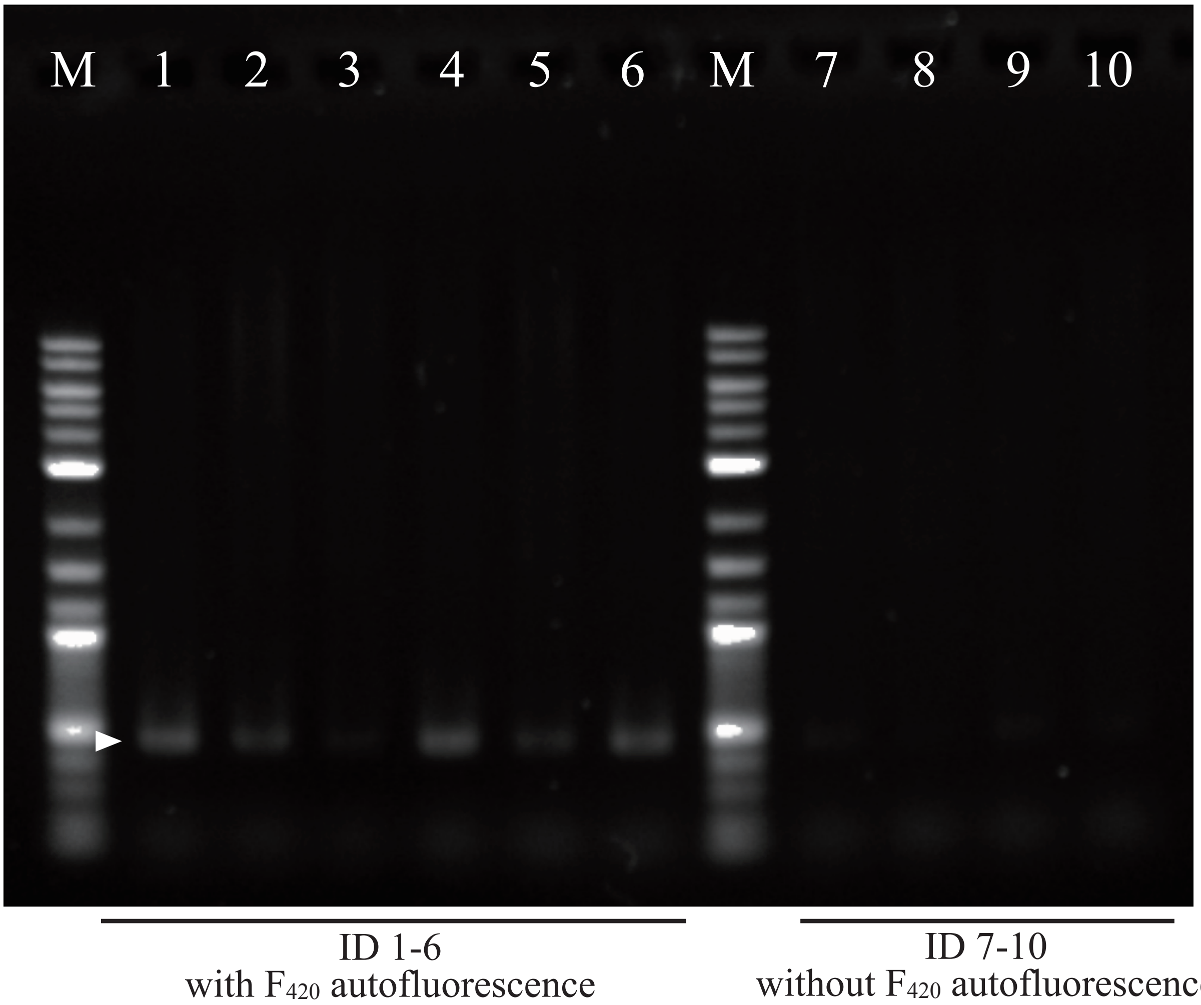


(B)

| Lane | 1 | 2 | 3 | 4 | 5 | 6 | 7 | 8 | 9 | 10 |
|---|---|---|---|---|---|---|---|---|---|---|
| Colony ID | O2022a | K2023a | K2023b | K2023e | O2023a | O2023e | O2022b | O2023b | O2023c | O2023d |
| Infection rate (%) | 98.2 | 97.6 | 94.7 | 99.7 | 91.5 | 84.1 | 0.0 | 3.9 | 0.6 | 0.2 |
| Calculated DNA conc. (µg/µL) | 44.3 | 20.9 | 6.3 | 42.1 | 16.0 | 38.4 | n.d. | n.d. | n.d. | n.d. |
| Methane (nmol/g/h) | n.a. | 89.9 | 80.4 | 79.8 | n.a. | n.a. | n.a. | 13.2 | 17.8 | 11.4 |

**Fig. S2.** Quantification of PCR products of the *mcrA* gene obtained from the whole gut DNA of *Coptotermes formosanus* in different colonies. (A) Results of agarose gel electrophoresis. Arrowhead indicates the predicted size of the target *mcrA* region. (B) Infection rates of endosymbiotic methanogens, quantities of PCR products, and methane emission rates of worker individuals from each termite colony. Image Lab software v5.0 (Bio-Rad) was used for image capture and densitometric analysis. The DNA concentration was estimated by comparison with the Quick-Load purple low molecular weight DNA ladder marker (New England Labs). n.d., not detected; n.a., not analysed.

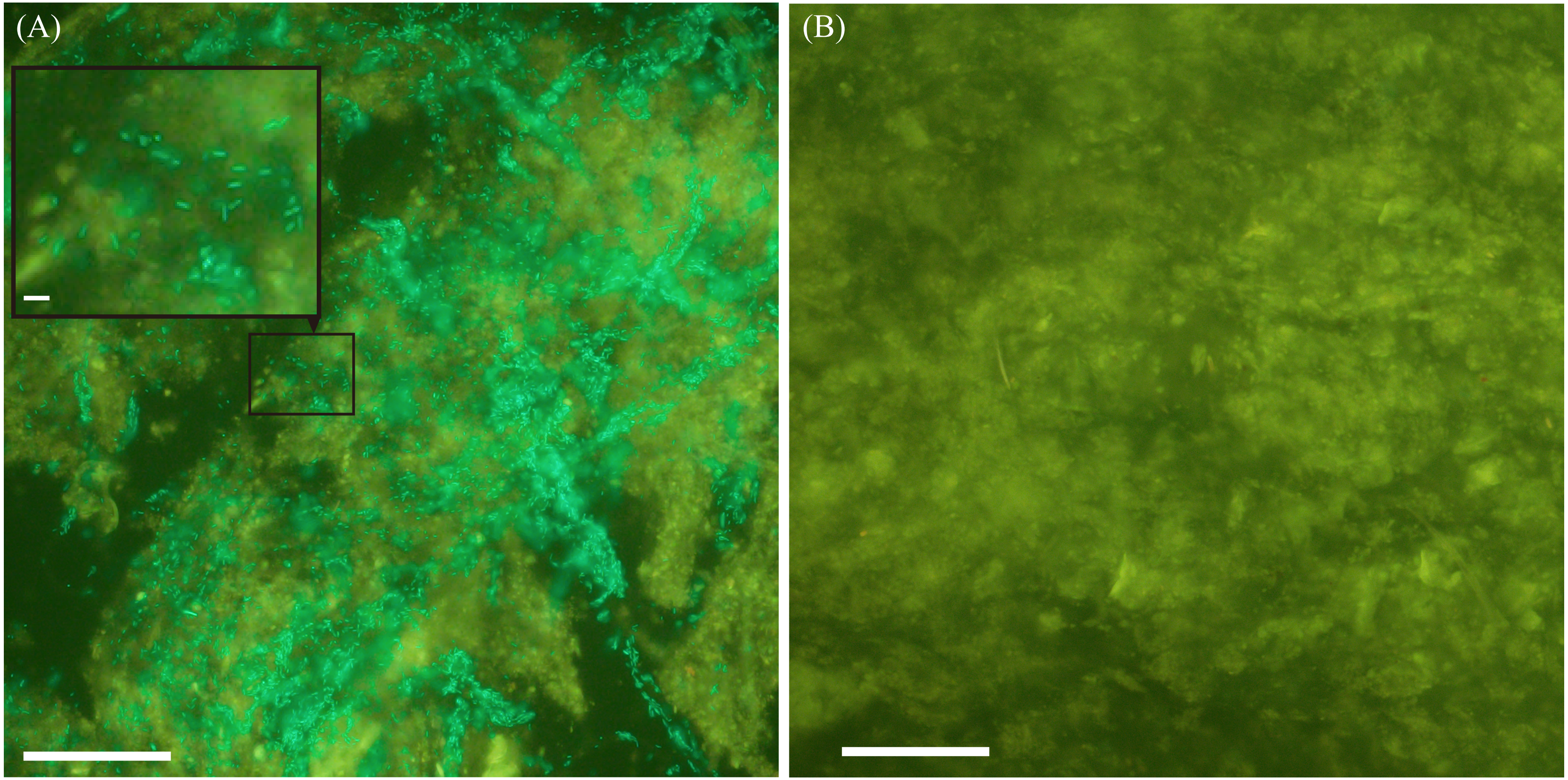


**Fig. S3** Detection of methanogens attached to the hindgut wall of *Coptotermes formosanus*. (A) Fragment of the hindgut wall of a worker termite from colony K2023c in which most *Cononympha* cells harboured methanogens (Table S2). Rod-shaped methanogens were visible with greenish autofluorescence of cofactor $F_{420}$. (B) Fragment of the hindgut wall of a worker termite from colony O2023b in which most *Cononympha* cells were without methanogens (Table S1). No $F_{420}$ autofluorescence was detected. Amorphous yellow autofluorescence was emitted by the termite tissues. Scale bars: 50 µm; inset of (A) = 1 µm.

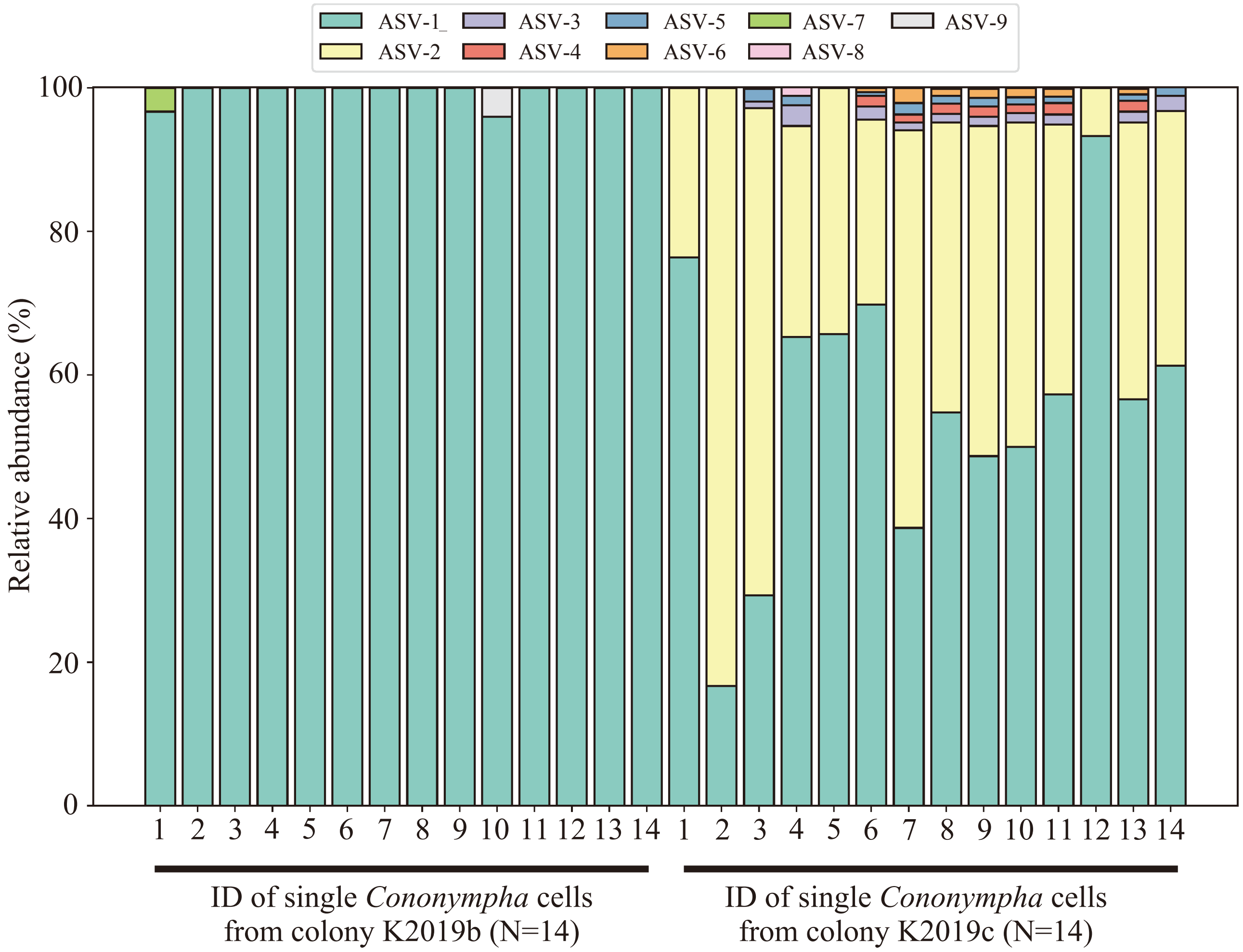


**Fig. S4.** Composition of 16S rRNA amplicon sequence variants (ASVs) assigned to *Methanobrevibacter* from single *Crononympha* cells. The most dominant ASV-1 and ASV-2 differed by one of 386 bases.

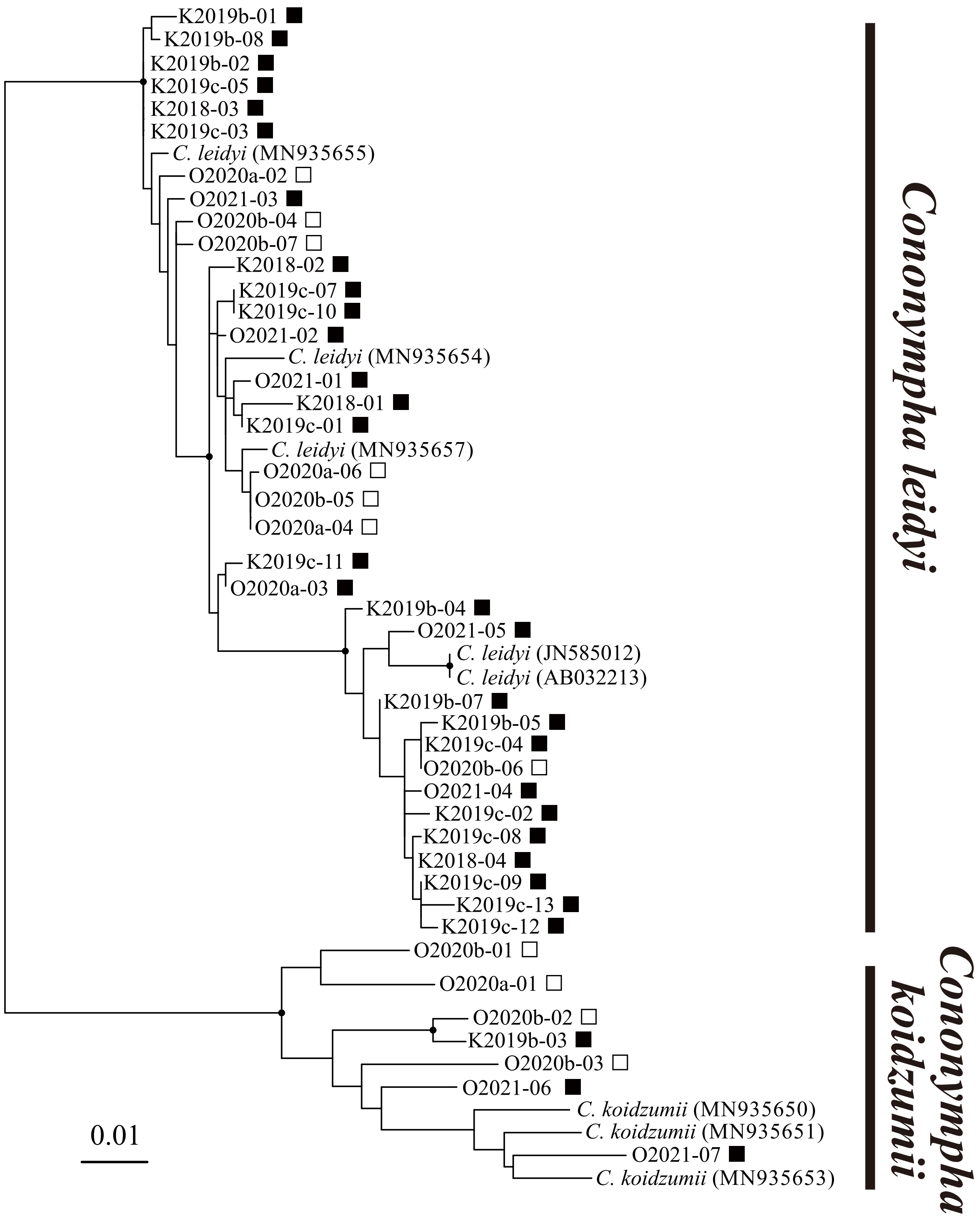


**Fig. S5.** Phylogenetic relationships of single *Cononympha* cells and identification of species (*i.e.*, *C. leidyi* or *C. koidzumii*) based on the 18S rRNA gene. Maximum likelihood tree was constructed using unambiguously aligned 814 nucleotide sites. Bootstrap values (>95%) with 1,000 replicates are shown as closed circles. *Cononympha* cells with endosymbiotic methanogens are shown with filled squares and those without endosymbiotic methanogens are shown with open squares. *Cononympha* cell IDs correspond to those in Figure S1, Figure 2, and Table S5. Scale bar indicates nucleotide substitutions per site.

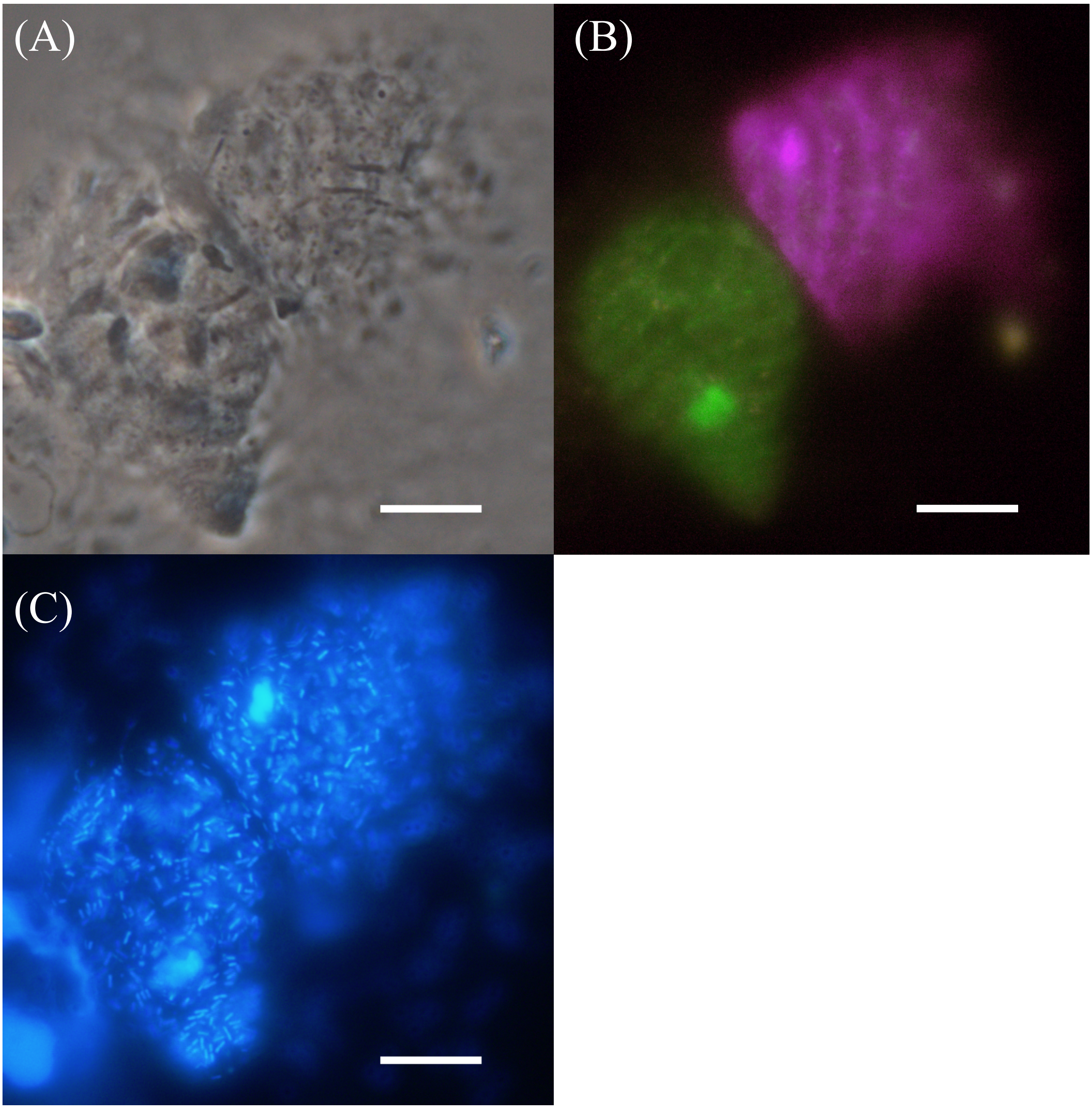


**Fig. S6** Detection of *Methanobrevibacter*-like rods in *Cononympha leidyi* and *Cononympha koidzumii* cells. (A) Phase-contrast image of two *Cononympha* cells. (B) Merged image of fluorescence in situ hybridization specifically targeting 18S RNA of *C. leidyi* (Texas red, magenta) and *C. koidzumii* (6FAM, green). Probes CLe-18S-177 and CKo-18S-177 in Table S1 were used. The original colour of Texas red was converted to magenta using Adobe Photoshop. (C) 4,6-Diamidino-2-phenylindole-stained DNA (blue). Bars = 10 µm.

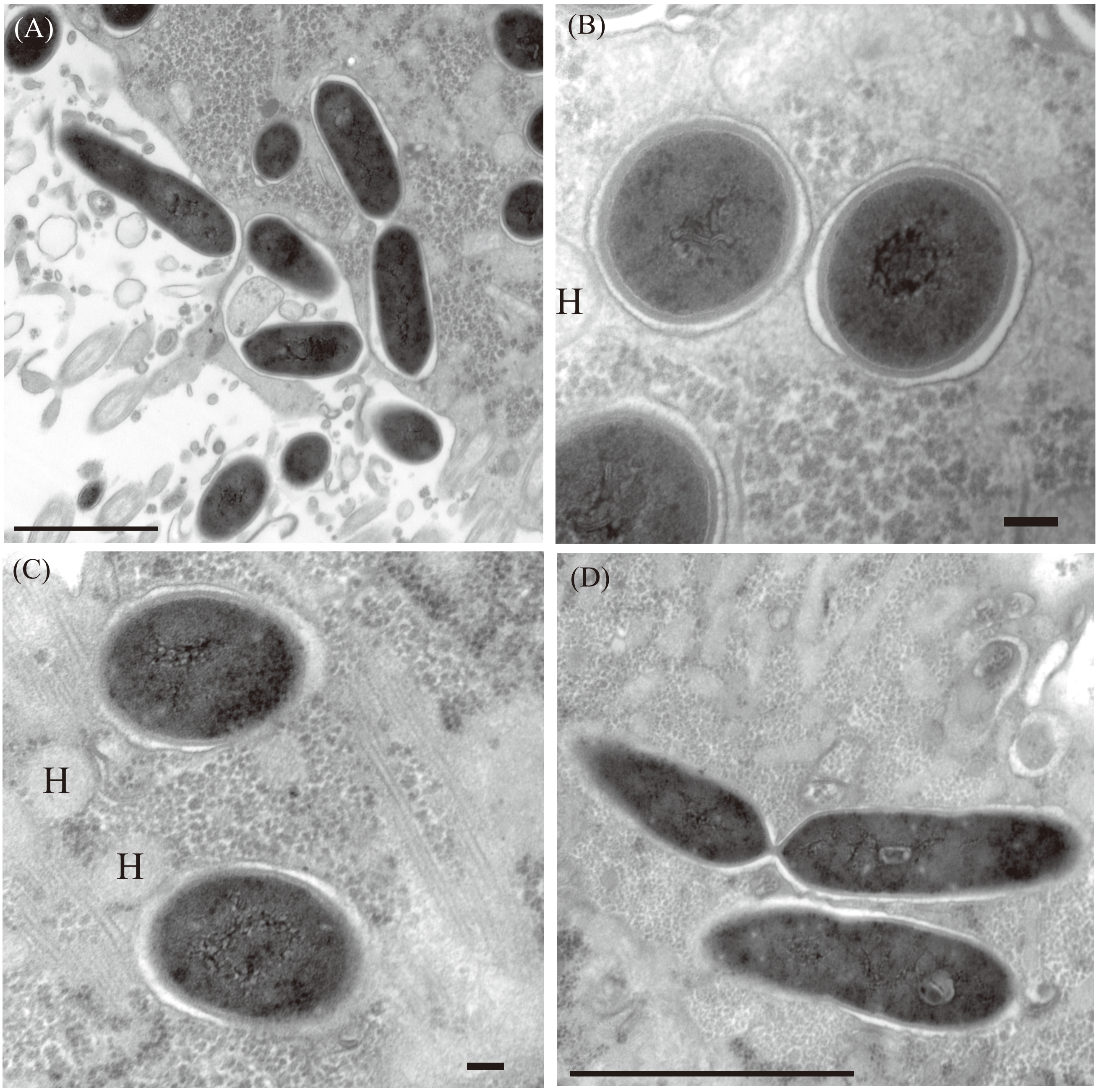


**Fig. S7** Transmission electron microscopy of *Cononympha* cells. (A–C) Magnified views of several parts of the *Cononympha* cell shown in Figure 3A. Electron-dense, *Methanobrevibacter*-like cells were observed. Each *Methanobrevibacter*-like cell was surrounded by an apparently host-derived membrane. (D) *Methanobrevibacter*-like cell during fission in another *Cononympha* cell. H: examples of putative hydrogenosomes. Bars = 1 μm (A, D); 100 nm (B, C).

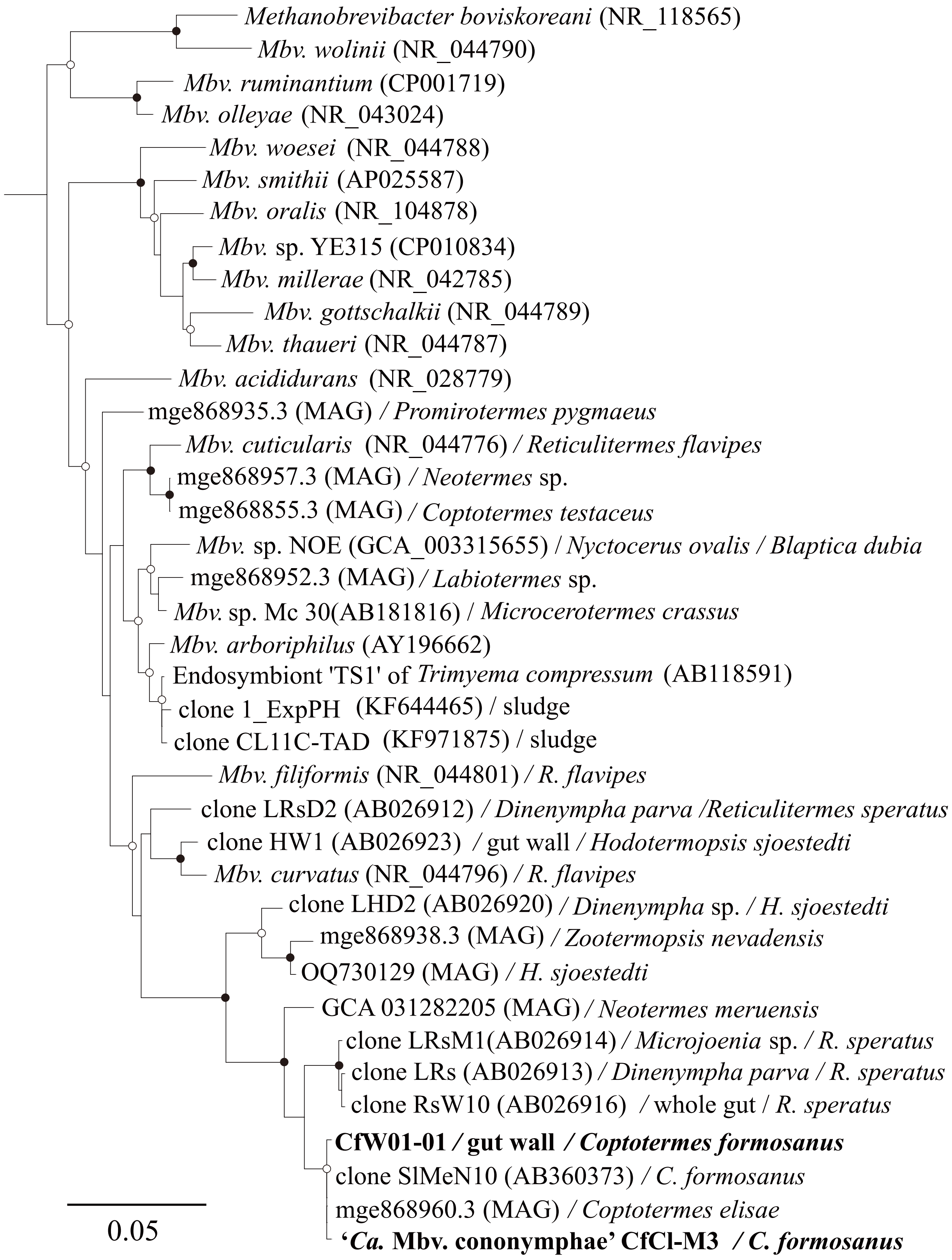


**Fig. S8.** Phylogenetic position of '*Ca.* Methanobrevibacter cononymphae' CfCl-M3 based on the 16S rRNA gene. Maximum likelihood tree was constructed using 1,408 unambiguously aligned nucleotide sites and rooted with *Methanothermus fervidus* (NR102926), *Methanobacterium formicicum* (NR025028), *Methanobacterium bryantii* (NR042781), and *Methanosphaera cuniculi* (NR104874) as outgroups. The sources of sequences or isolates are shown (e.g., protist species/termite species). Bootstrap support values with 1,000 replicates are indicated with closed circles (>95%) and open circles (95–80%). Scale bar indicates nucleotide substitutions per site.

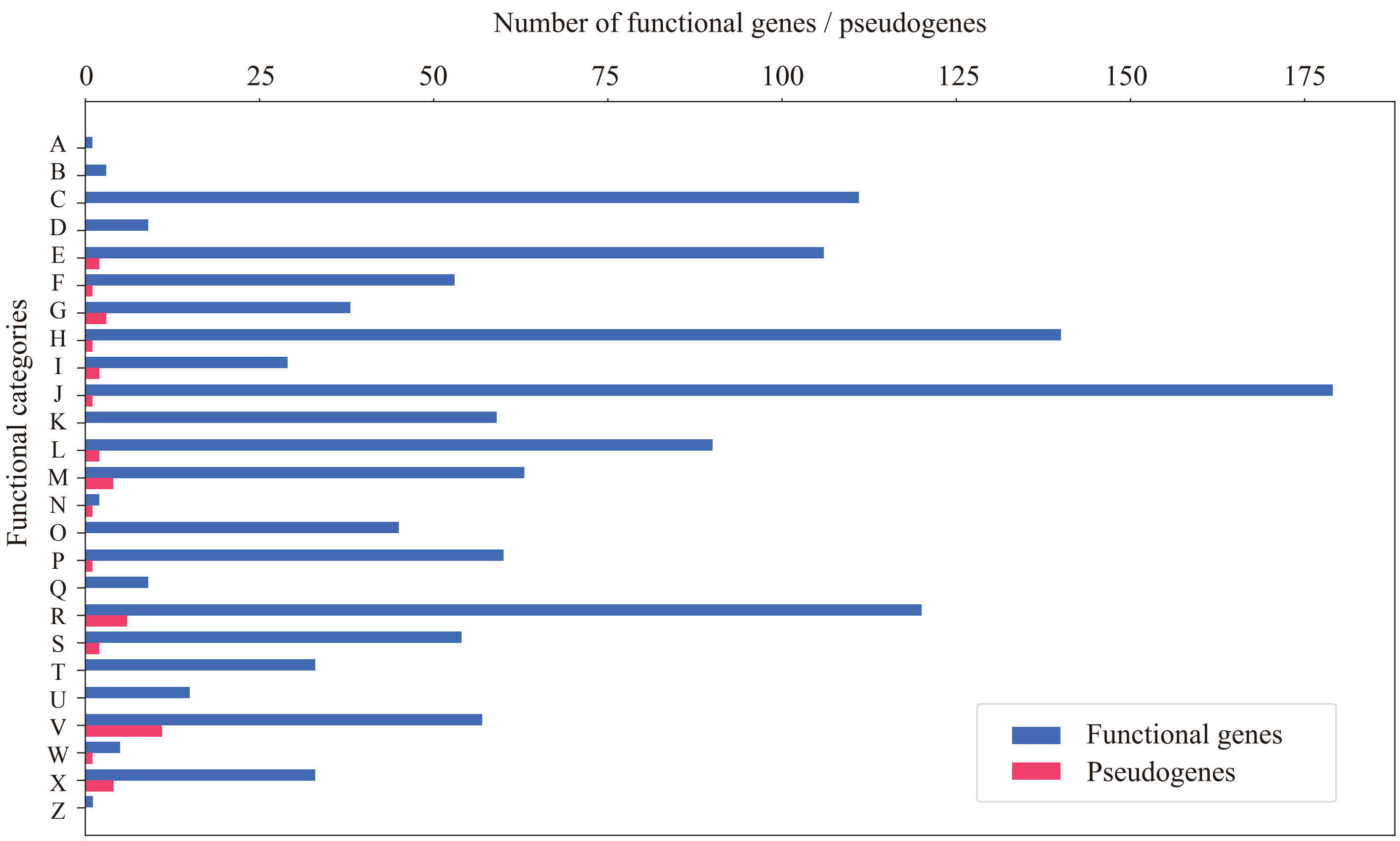


**Fig. S9.** Functional classification of genes and pseudogenes of '*Ca.* Methanobrevibacter cononymphae' CfCl-M3. Categories according to the Clusters of Orthologous Genes (COG) database (Tatusov et al., 1997) are as follows: (A) RNA processing and modification; (B) chromatin structure and dynamics; (C) energy production and conversion; (D) cell cycle control, cell division, chromosome partitioning; (E) amino acid transport and metabolism; (F) nucleotide transport and metabolism; (G) carbohydrate transport and metabolism; (H) coenzyme transport and metabolism; (I) lipid transport and metabolism; (J) translation, ribosomal structure, and biogenesis; (K) transcription; (L) replication, recombination, and repair; (M) cell wall/membrane/envelope biogenesis; (N) cell motility; (O) posttranslational modification, protein turnover, and chaperones; (P) inorganic ion transport and metabolism; (Q) secondary metabolite biosynthesis, transport, and catabolism; (R) general function prediction only; (S) function unknown; (T) signal transduction mechanisms; (U) intracellular trafficking, secretion, and vesicular transport; (V) defence mechanisms; (W) extracellular structures; (X) mobilome: prophages, transposons; (Z) cytoskeleton.

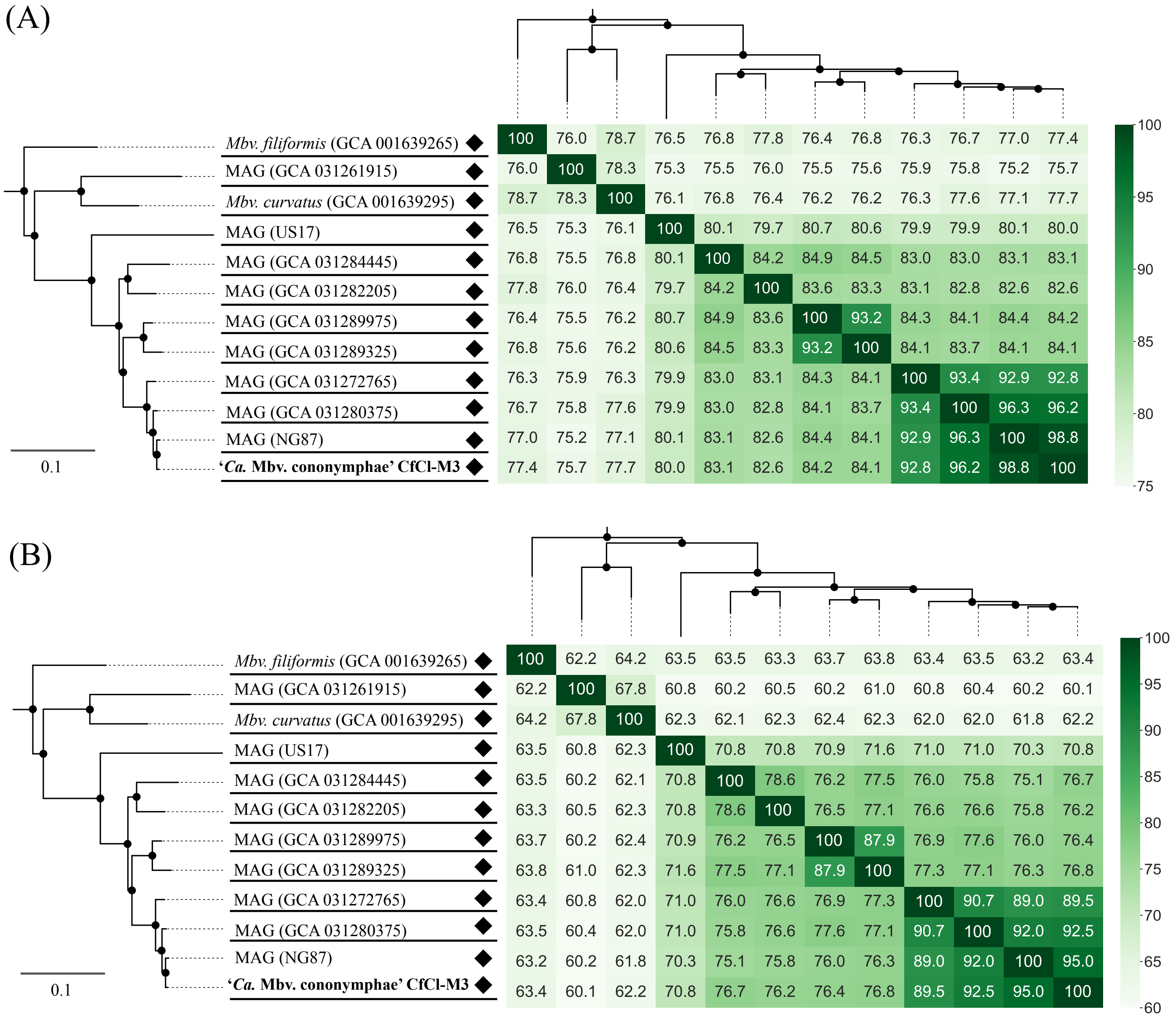


**Fig. S10.** Pairwise ANI (A) and AAI (B) heatmaps of the genomes in the clade including '*Ca.* Methanobrevibacter cononymphae'.

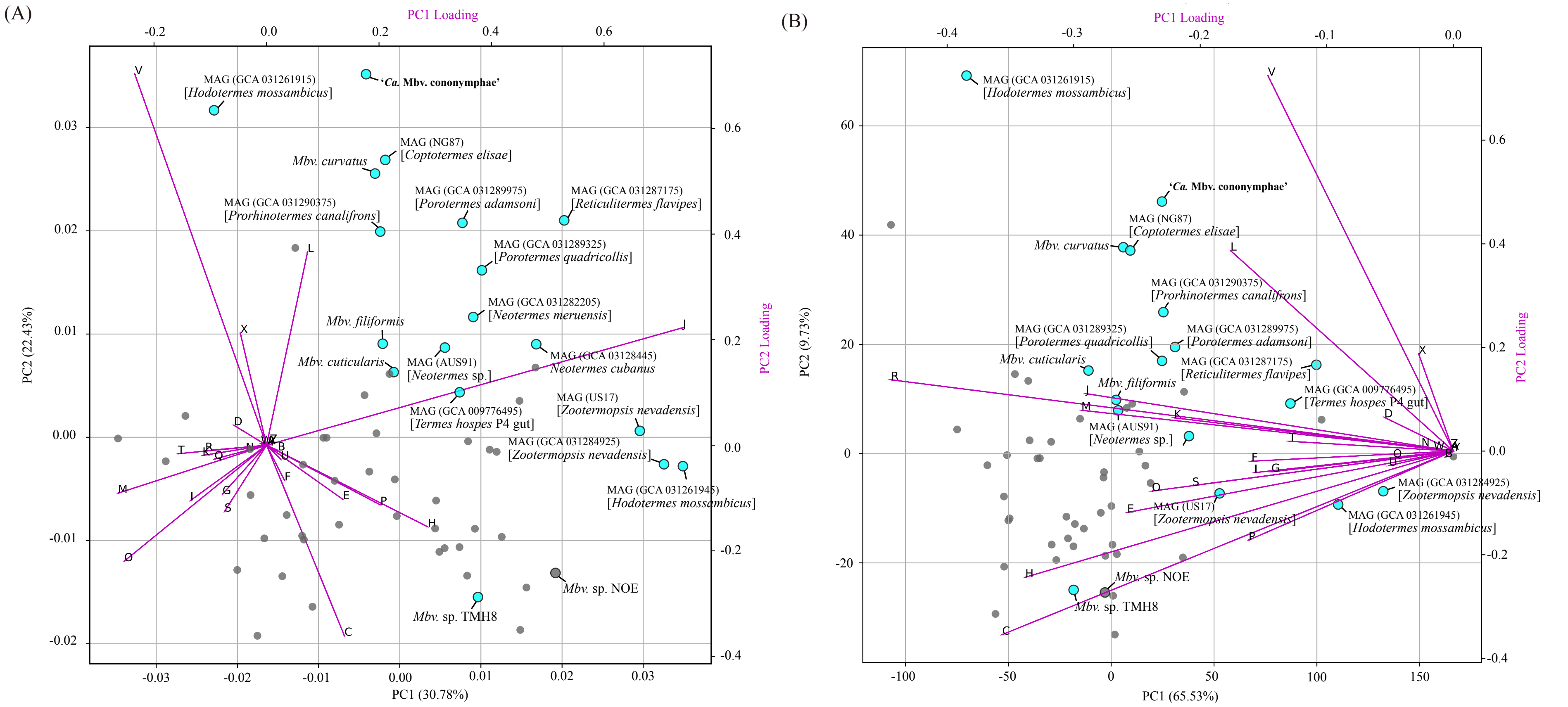


**Fig. S11.** Principal component analysis of *Methanobrevibacter* genomes based on the relative abundance (A) and the number (B) of genes assigned to respective COG (clusters of orthologous genes) functional categories. Genome sequences derived from termite guts are highlighted in cyan.

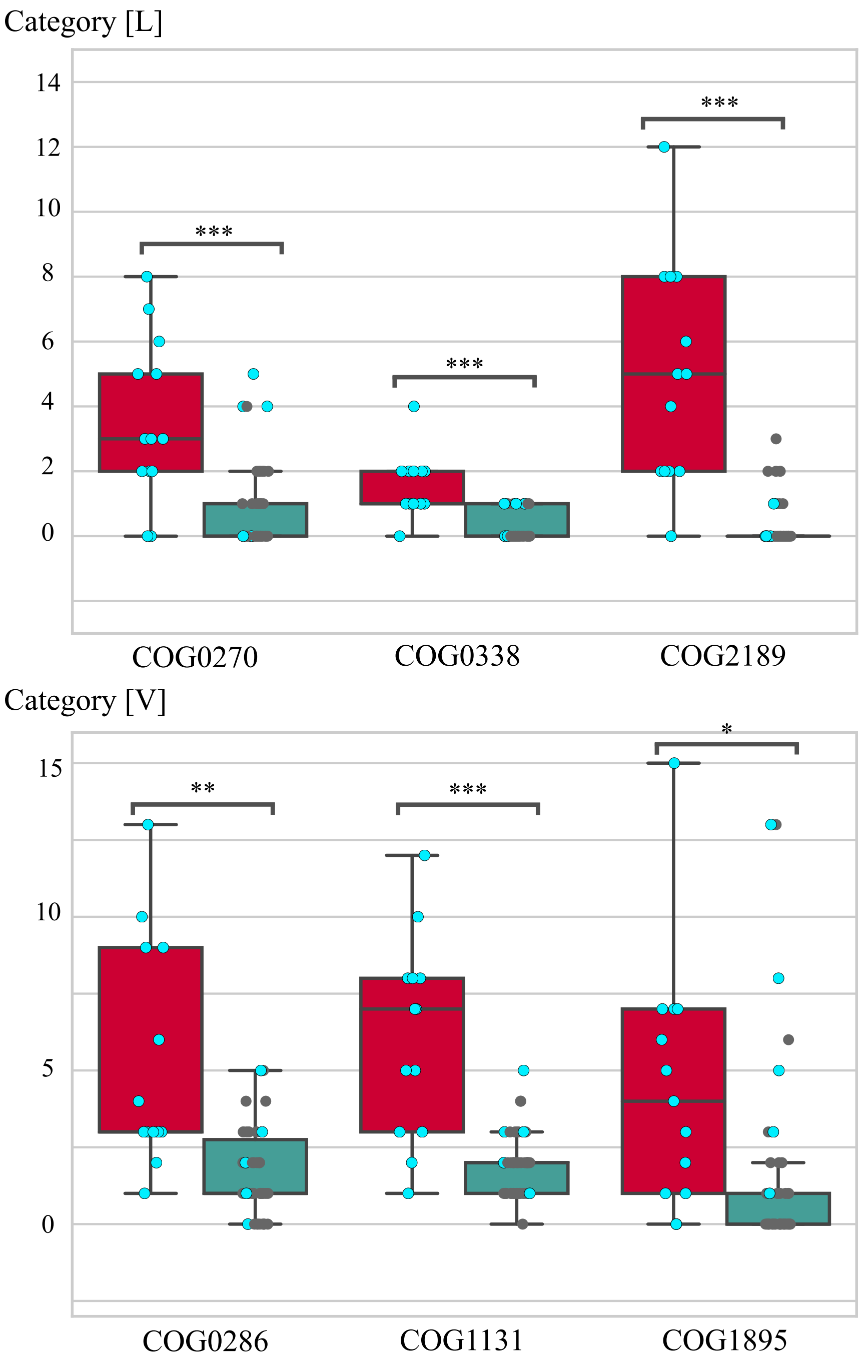


**Fig. S12.** Comparison of the number of genes assigned to COG (clusters of orthologous genes) functional categories [L] and [V] between clades of *Methanobrevibacter*. A clade containing '*Ca.* Methanobrevibacter cononymphae' (red) and its sister clade (green) in Fig. 4 were compared. Cyan dots represent the genome sequences derived from termite guts. Two functional categories (L: replication, recombination, and repair; V: defence mechanisms) were selected because they chiefly contributed to the position of *Mbv. cononymphae* in the PCA plot in Fig. S11. COG0270 corresponds to cytosine methylase; COG0338, adenine methylase; COG2189, adenine methylase Mod; COG0286, methylase subunit of Type I restriction-modification system; COG1131, ATPase component of ABC-type multidrug transport system; COG1895, HEPN domain protein of predicted MNT-HEPN system toxin. Student's *t*-test was performed: *** <0.001; ** <0.01; * <0.05.

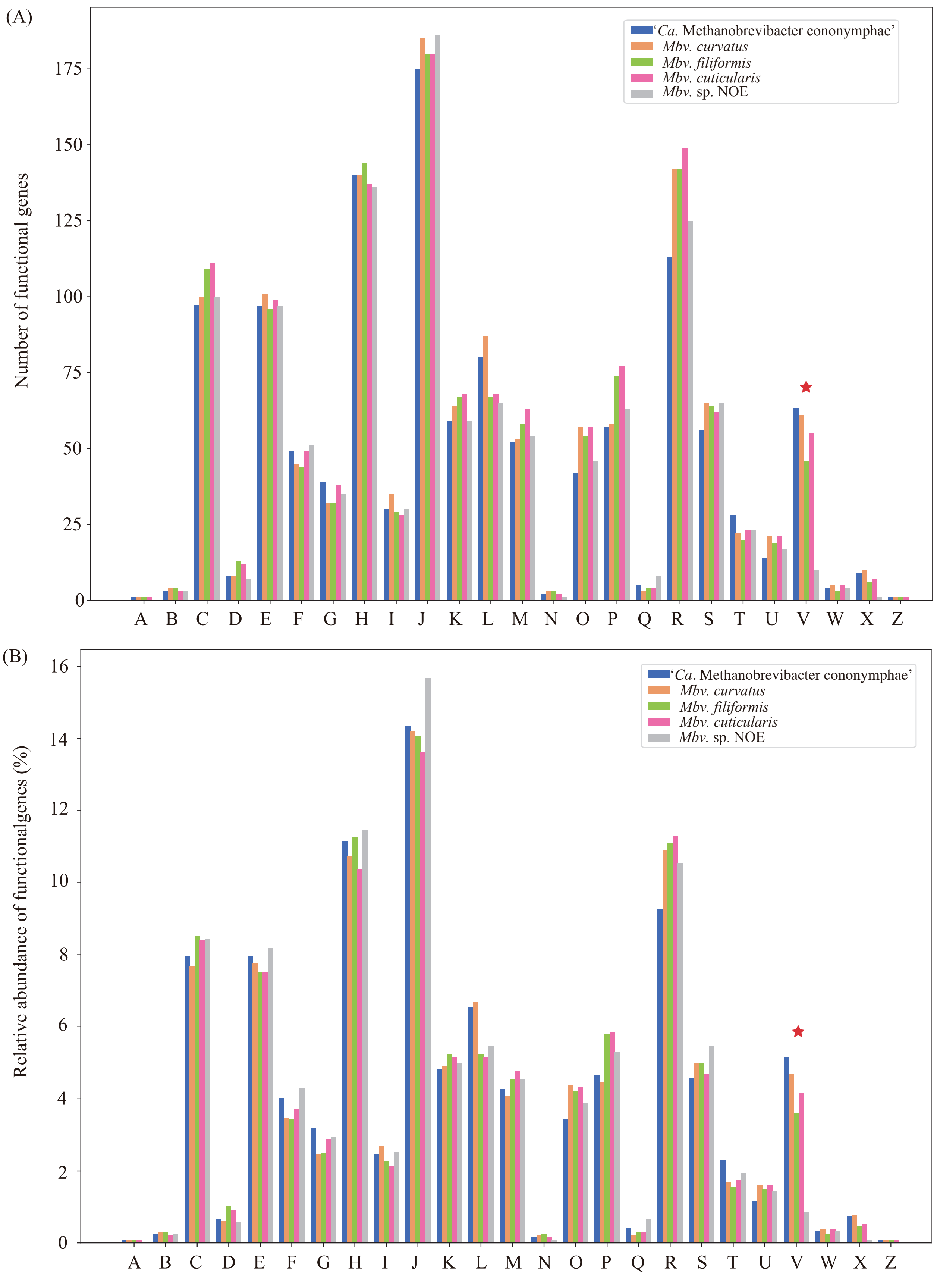


**Fig. S13.** Comparison of the number (A) and ratio (B) of genes classified into COG (clusters of orthologous genes) functional categories. The genome of '*Ca.* Methanobrevibacter cononymphae' was compared with three *Methanobrevibacter* species isolated from the guts of *Reticulitermes flavipes* and the methanogenic endosymbiont NOE of the ciliate *Nyctotherus ovalis*. Abbreviations of COG categories are explained in the legend to Figure S9.

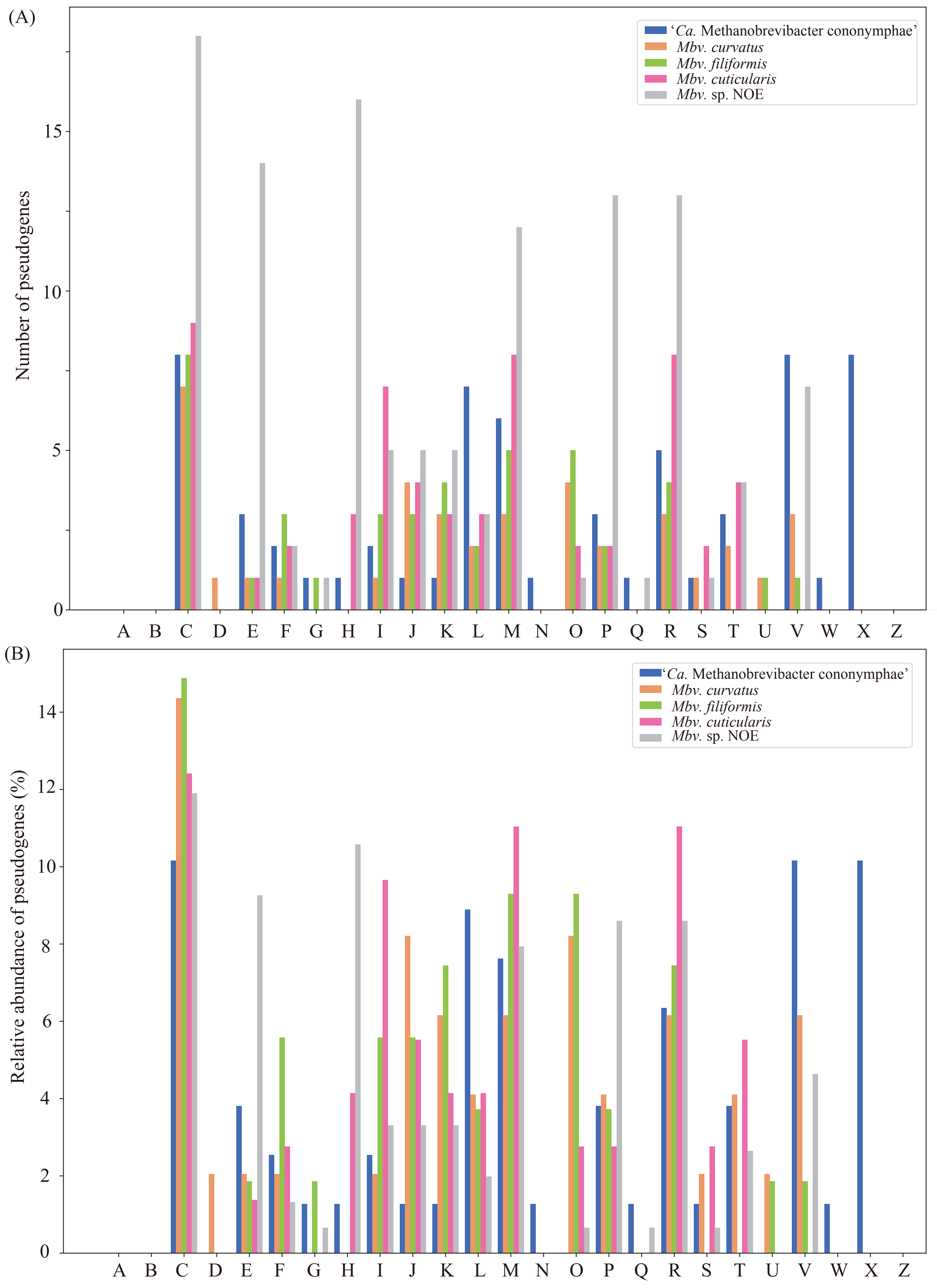


**Fig. S14.** Comparison of the number (A) and the ratio (B) of pseudogenes classified into COG (clusters of orthologous genes) functional categories. See also the legend to Figure S13.

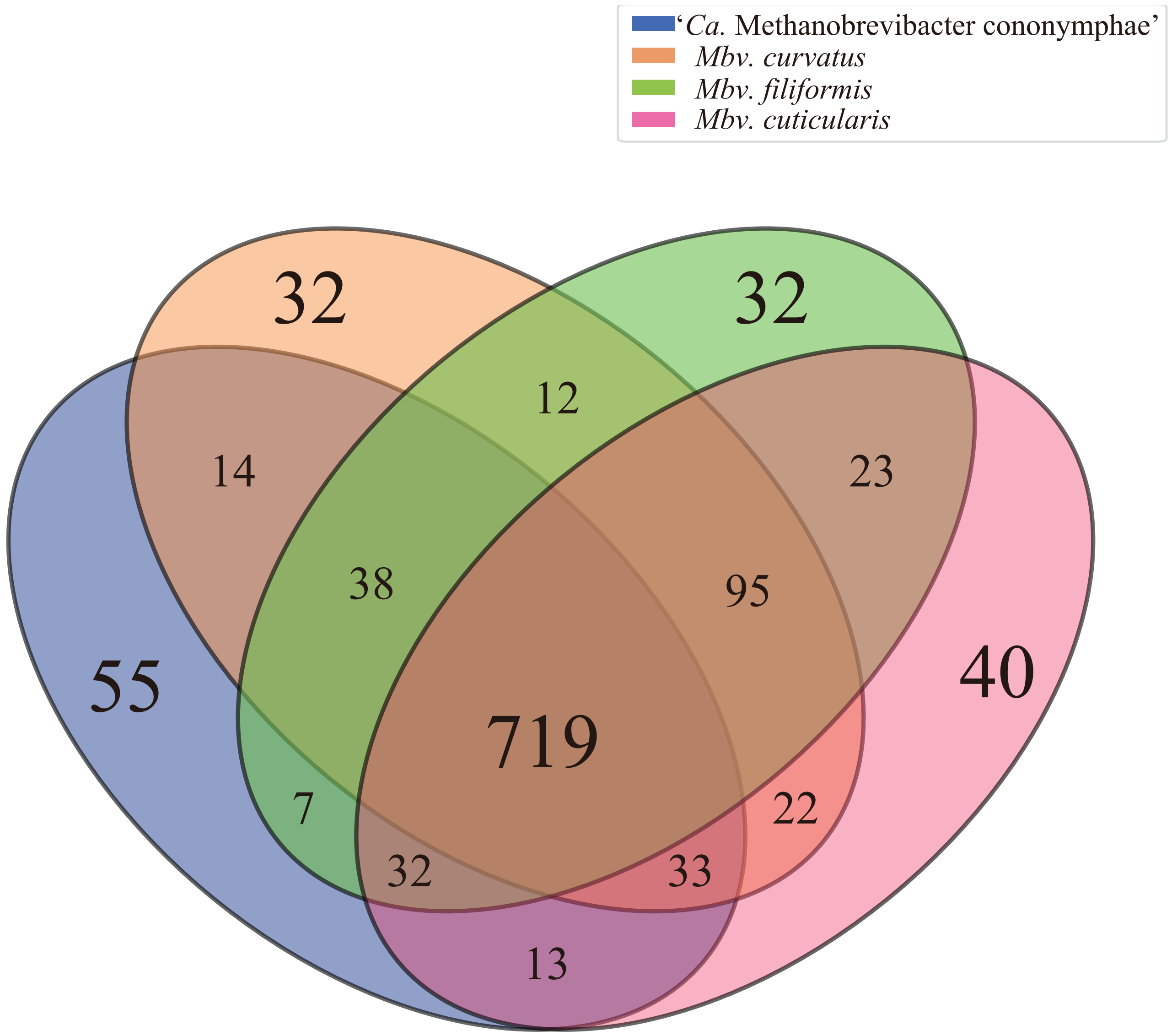


**Fig. S15.** Venn diagram of genes classified into clusters of orthologous genes. The genome of '*Ca.* Methanobrevibacter cononymphae' was compared with those of three *Methanobrevibacter* species isolated from the guts of *Reticulitermes flavipes* (Poehleina and Seedorf, 2016). Details of the unique genes of '*Ca.* Methanobrevibacter cononymphae' are shown in Table S10.

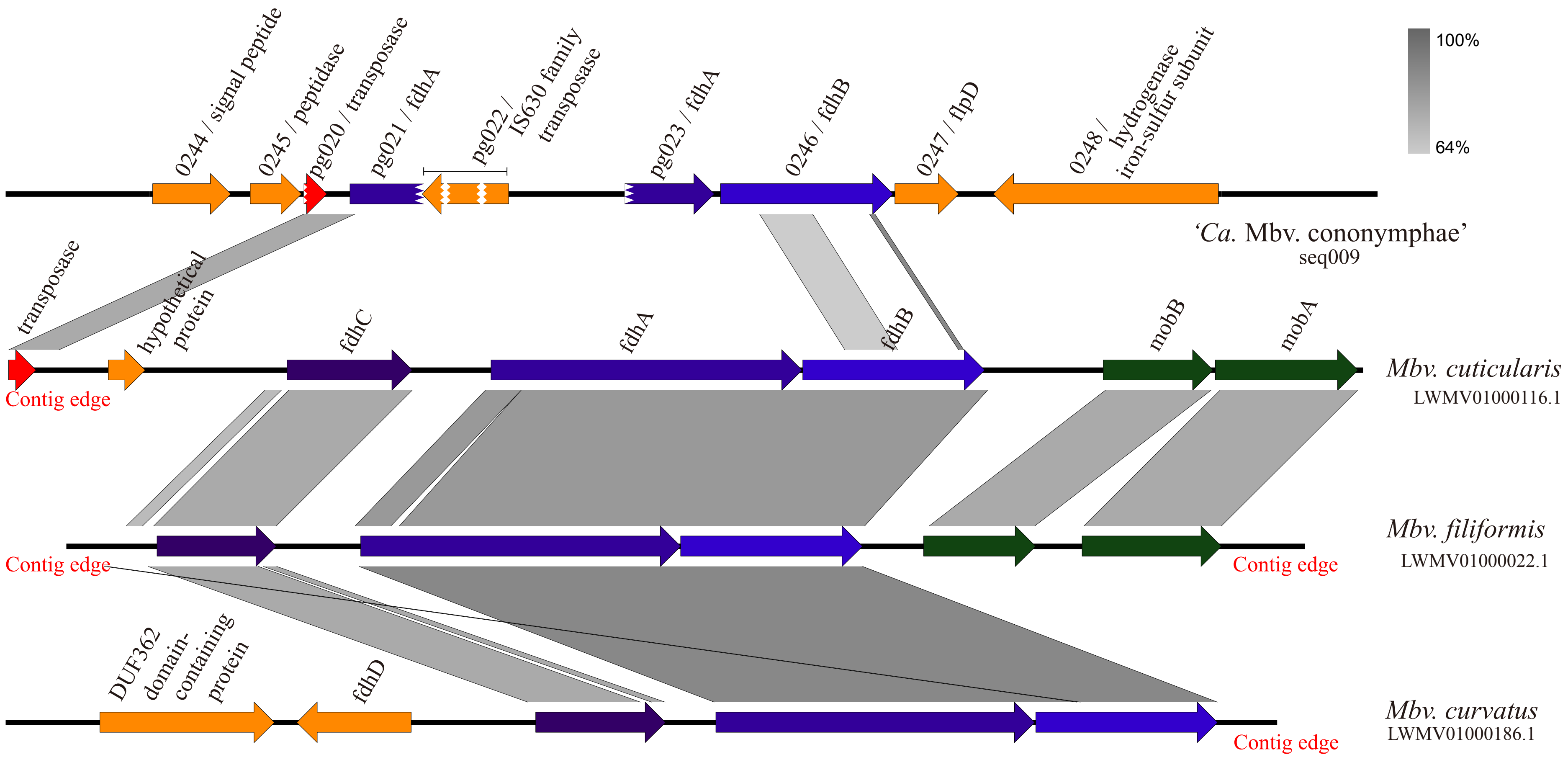


**Fig. S16.** Comparison of gene clusters encoding formate dehydrogenase FdhAB and formate transporter FdhC. Truncated arrows represent pseudogenes. Genes *mobA* (cyclic pyranopterin monophosphate synthase), *mobB* (molybdoterin-guanine dinucleotide biosynthesis protein), and *fdhD* (formate dehydrogenase accessory sulfurtransferase) are encoded in other contigs of '*Ca.* Methanobrevibacter cononymphae'. Gene IDs correspond to those in Table S4.
